# p73 activates transcriptional signatures of basal lineage identity and ciliogenesis in pancreatic ductal adenocarcinoma

**DOI:** 10.1101/2023.04.20.537667

**Authors:** Stella K. Hur, Tim D.D. Somerville, Xiaoli S. Wu, Diogo Maia-Silva, Osama E. Demerdash, David A. Tuveson, Faiyaz Notta, Christopher R. Vakoc

**Affiliations:** Cold Spring Harbor Laboratory, Cold Spring Harbor, NY 11724, U.S.A; Department of Medical Biophysics, University of Toronto, Toronto, Ontario, Canada

## Abstract

During the progression of pancreatic ductal adenocarcinoma (PDAC), tumor cells are known to acquire transcriptional and morphological properties of the basal (also known as squamous) epithelial lineage, which leads to more aggressive disease characteristics. Here, we show that a subset of basal-like PDAC tumors aberrantly express p73 (TA isoform), which is a known transcriptional activator of basal lineage identity, ciliogenesis, and tumor suppression in normal tissue development. Using gain- and loss- of function experiments, we show that p73 is necessary and sufficient to activate genes related to basal identity (e.g. *KRT5*), ciliogenesis (e.g. *FOXJ1*), and p53-like tumor suppression (e.g. *CDKN1A*) in human PDAC models. Owing to the paradoxical combination of oncogenic and tumor suppressive outputs of this transcription factor, we propose that PDAC cells express a low level of p73 that is optimal for promoting lineage plasticity without severe impairment of cell proliferation. Collectively, our study reinforces how PDAC cells exploit master regulators of the basal epithelial lineage during disease progression.

## Introduction

Pancreatic ductal adenocarcinoma (PDAC) is one of the most lethal human cancers due to lack of early detection methods and limited treatment options (Siegel & Miller, 2019). Extensive genetic, transcriptomic, and histological studies have illustrated that inter- and intra-tumoral heterogeneity exists in PDAC, which poses significant treatment obstacles. This issue motivated several studies that seek to molecularly classify tumors to provide better insights into disease biology (Bailey et al., 2016; Chan-Seng-Yue et al., 2020; Collisson et al., 2011; Hayashi et al., 2020; Moffitt et al., 2015; The Cancer Genome Atlas Research Network, 2017). Transcriptome studies identified two major transcriptional subtypes of PDAC: classical subtype (also known as the progenitor subtype) that expresses pancreatic endoderm specification genes such as *GATA6*, *HNF1B*, *HNF4A*, and *PDX1*, and the basal-like subtype (also known as squamous subtype) that expresses low levels of the endoderm markers and high levels of basal/squamous lineage genes such as *KRT5*, *KRT6A*, *S100A2*, *PTHLH*, and *TP63* (Bailey et al., 2016; Collisson et al., 2011; Moffitt et al., 2015; Somerville et al., 2018; The Cancer Genome Atlas Research Network, 2017). Notably, basal-like PDAC is associated with a shorter survival than classical PDAC (Bailey et al., 2016; Chan-Seng-Yue et al., 2020; Collisson et al., 2011; Hayashi et al., 2020; Moffitt et al., 2015; Somerville et al., 2018; The Cancer Genome Atlas Research Network, 2017).

Subsequent studies demonstrated that transcriptional programs that define these subtypes are responsive to various cell intrinsic or extrinsic stimuli, which can result in switching of tumor subtypes (Miyabayashi et al., 2020; Somerville et al., 2018; Somerville et al., 2020). For example, we have shown that transcription factor p63 (encoded by the *TP63* gene) plays a causal role in establishing and maintaining the basal/squamous lineage program by remodeling the enhancer landscape (Somerville et al., 2018). In addition, we have shown that ZBED2 is preferentially expressed in basal-like PDAC and inhibits expression of *GATA6*, a master regulator of classical PDAC (Somerville et al., 2020). Notably, both p63 and ZBED2 are selectively expressed in the normal basal/squamous epithelial lineage in addition to their acquired expression in basal-like PDAC.

p73 (encoded by the *TP73* gene) belongs to the p53 family of transcription factors and is expressed from two different promoters to produce transcripts that either retain (referred to hereafter as *TA*) or exclude (referred to hereafter as *dN*) an N-terminal transactivation domain (Kaghad et al., 1997; Yang et al., 2000). In addition, alternative splicing of the C-terminus of *TP73* has been described (Kaghad et al., 1997; Laurenzi et al., 1998; Laurenzi et al., 1999). It is thought that p73-TA activates p53-like responses such as apoptosis and cell cycle arrest (Melino et al., 2002; Zhu et al., 1998), whereas p73-dN exhibits oncogenic behaviors (Grob et al., 2001). Importantly, expression of p73 isoforms is tissue-specific, therefore requires careful validation of their protein expression and function in the corresponding cell types (Grespi et al., 2012; Vikhreva et al., 2018).

Mice deficient in p73 (*Trp73* is the mouse gene symbol) have provided insights into various developmental roles of this transcription factor, including regulation of neurodevelopment, fertility, and inflammation (Tomasini et al., 2008; Wilhelm et al., 2010; Yang et al., 2000). Notably, recent reports have discovered that much of the total p73 knock-out mouse phenotypes can be explained by loss of multiciliated cells, thus implicating p73-TA as a master regulator of ciliogenesis (Marshall et al., 2016; Nemajerova et al., 2016). Notably, p73 total knock-out mice display more severe developmental defects than the individual p73-TA or p73-dN isoform-specific knock-out mice (Tomasini et al., 2008; Wilhelm et al., 2010; Yang et al., 2000). Therefore, it is possible that despite the seemingly antagonistic function of TA and dN isoform mentioned above (Grob et al., 2001), these isoforms may intricately collaborate or play redundant role during development (Tomasini et al., 2008; Wilhelm et al., 2010; Yang et al., 2000).

p73 overexpression has been detected in several different cancer lineages (Deyoung & Ellisen, 2007; Dominguez et al., 2001; Niklison-chirou et al., 2017; Rufini et al., 2011; Yokomizo et al., 1999). However, it has been challenging to define a role of p73 in tumorigenesis. Studies that measured relative p73 isoform expression in various types of tumors found that increase in the ratio of dN:TA or overexpression of p73-dN itself generally correlates with disease progression (Rufini et al., 2011). Yet, p73-TA is expressed at a higher level than p73-dN in a subset of cancer cell lines (Conforti et al., 2012); p73-TA sustained proliferation of medulloblastoma cells by regulating glutamine metabolism (Niklison-chirou et al., 2017); p73-TA induced expression of pro-inflammatory cytokine interleukin-1β in non-small cell lung cancer cells, suggestive of tumor-promoting role (Vikhreva et al., 2017).

Interestingly, total p73 knock-out mice lacked spontaneous tumor formation, while p73-TA knock-out mice were more prone to spontaneous tumorigenesis, adding to the view that p73-TA is a tumor suppressor (Tomasini et al., 2008; Yang et al., 2000). Collectively, a complex relationship exists between p73 isoforms and normal and neoplastic developmental process.

While p53 and p63 have well-described roles in PDAC biology, there is limited description of p73 function in the context of PDAC. In this study, we show that p73-TA is expressed in basal-like human PDAC and correlates with a poor prognosis. Experimentally, we show that moderate expression of p73-TA can activate the basal lineage and ciliogenesis transcriptional programs and induces cell migration and invasion. These findings provide insights into the previously unrecognized dosage-dependent roles of p73 in PDAC biology.

## Results

### TP73-TA expression correlates with inferior prognosis and is preferentially expressed in basal-like PDAC

p63 and p73 are known to cooperate in the development of normal basal epithelial cells (Marshall et al., 2016). Since PDAC cells are known to acquire p63 expression to promote basal lineage identity in this tumor, we considered whether p73 was also involved in this cancer context. We began by characterizing the expression pattern of *TP73* in published transcriptome from human PDAC. In three independent clinical cohorts, we noted that patients with tumors expressing higher levels of *TP73* (*TP73*^high^) presented shorter overall survival compared to those with tumors expressing lower levels of *TP73* (*TP73*^low^) (Fig. 1 A, Supplementary Figure 1 A) (Bailey et al., 2016; Moffitt et al., 2015; The Cancer Genome Atlas Research Network, 2017). Although there was a minimal difference in tumor stage between the two groups, *TP73*^high^ tumors were more frequently classified as poorly differentiated than the *TP73*^low^ tumors (Supplementary Figure 2 A). In addition, *TP73*^high^ tumors displayed more frequent mutations in *TP53* compared to the *TP73*^low^ tumors (Supplementary Figure 2 A). We validated that *TP73* was aberrantly expressed in organoid cultures derived from human PDAC tumors, whereas *TP73* was lowly expressed in organoids derived from healthy normal pancreas (Fig. 1 B) (Tiriac et al., 2018). Consistently, transcriptome analysis of normal tissues by the Genotype-Tissue Expression (GTEx) project revealed minimal expression of *TP73* in the normal human pancreas (Supplementary Figure 1 B) (The GTEx Consortium, 2015). We next checked whether *TP73* expression was skewed toward either of the previously identified transcriptional subtypes of PDAC. Specifically, given structural and functional similarities to *TP63* (Moll and Slade 2004), a known master regulator of the basal/squamous lineage (Mills et al., 1999; Soares & Zhou, 2018; Somerville et al., 2018), it was of particular interest whether *TP73* would be highly expressed in the more aggressive, basal-like subtype of PDAC. Remarkably, *TP73* was expressed at a higher level in the basal-like subtype compared to the progenitor subtype of PDAC (Fig. 1 C) (Bailey et al., 2016; Moffitt et al., 2015; The Cancer Genome Atlas Research Network, 2017). Collectively, these aggressive molecular and clinical features seen in patient tumor samples suggested that *TP73* may contribute to the disease progression.

**Figure 1.**
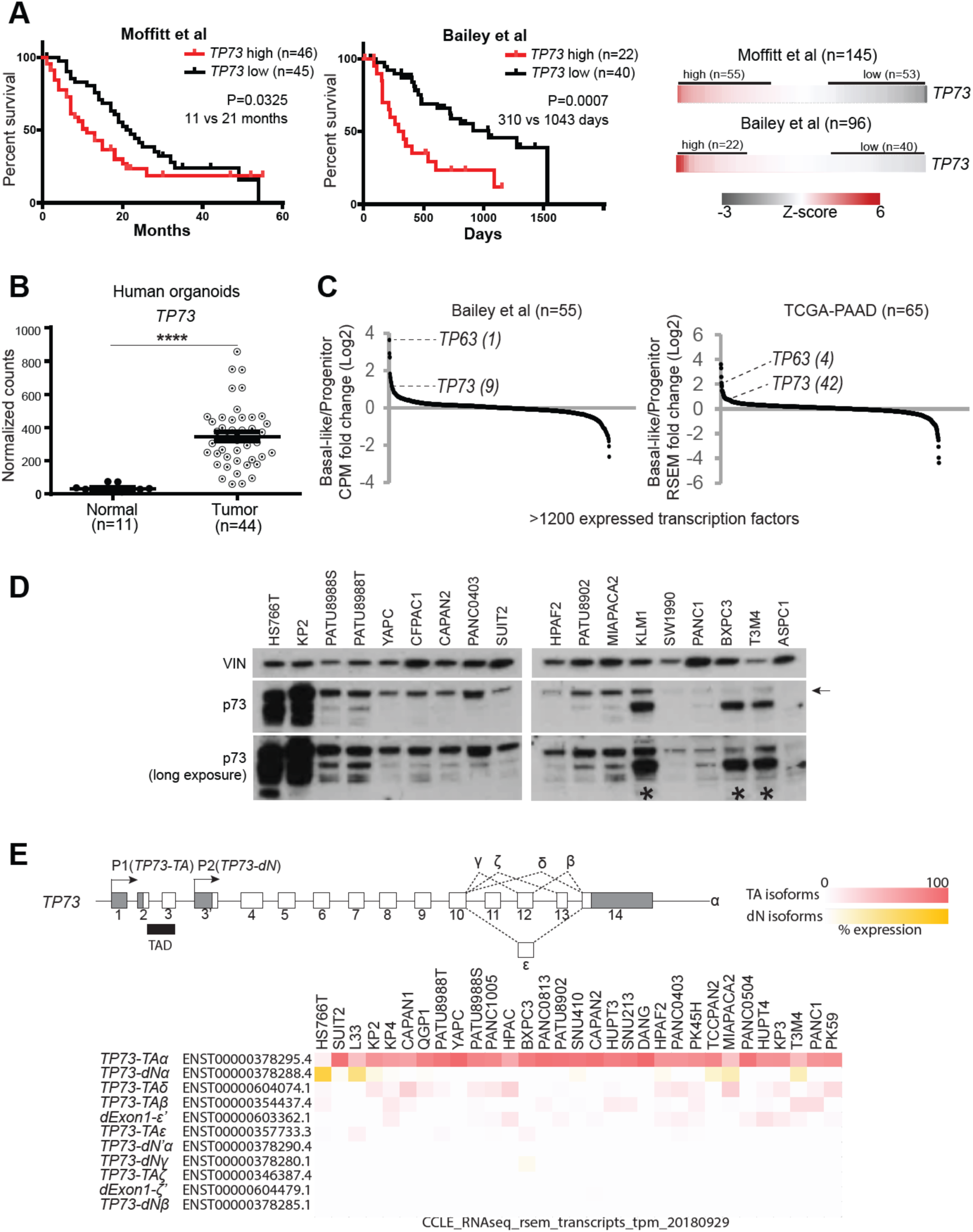
*TP73* is prognostic, aberrantly upregulated in the PDAC tumor organoids, and preferentially expressed in the squamous subtype of PDAC. (*A*) Survival curves of patients stratified by *TP73* expression. Log-rank (Mantel-Cox) test used to assess the median survival values and p-values. (*B*) Expression of *TP73* in human organoids derived from normal pancreata (Normal) or PDAC tumors (Tumor) (Tiriac et al., 2018). P-value calculated using unpaired t-test. ****, P ≤ 0.0001. Mean ± SEM shown. (*C*) Expression of transcription factors in the basal-like vs progenitor subtype of tumors. Transcription factors are ranked by their mean log2 fold-change in expression levels in the basal-like vs. progenitor subtype tumors. *TP63*, a master regulator of the squamous subtype PDAC (Somerville et al., 2018) is shown as a reference, and the rank of each gene is written inside parentheses. (*D*) Western blot analysis of TP73 in PDAC cell lines using a pan-TP73 antibody. VINCULIN (VIN) shown as a loading control. Bands corresponding to the molecular weight of the longest p73 isoform (i.e. p73-TAα) indicated with an arrow. Asterisk points to antibody cross-reactivity with p63-dN in cell lines expressing high levels of p63-dN. (*E*) (Top) Schematic of the human *TP73* gene. Exons (white boxes), UTRs (gray boxes), and introns (thin lines) are depicted. Exons are numbered. Transactivation domain (TAD) is indicated with black box. 3’ spliced isoforms are indicated in Greek letters. (Bottom) *TP73* isoform expression in PDAC cell lines from CCLE (Ghandi et al., 2019). Isoforms expressed from P1 or P2 promoter are referred to as *TP73-TA* or *TP73-dN*, respectively. Isoforms lacking exon1 but retaining the transactivation domain are noted as dExon1. 3’ spliced isoforms with variations at the 3’ UTR are marked with a prime (‘) symbol (i.e. dExon1-ε’ and dExon1-ζ’). Isoform expressed from an alternative promoter other than P1 or P2 is noted as dN’ (i.e. TP73- dN’α). For the heatmap, cell lines are listed in descending order (from left to right) of total *TP73* isoform expression. *TP73-TA* and *dExon1* are collectively considered as TA isoforms, and *TP73-dN* and *TP73- dN’* are collectively considered as dN isoforms. Expression of each isoform is normalized within a cell line (i.e. % expression = tpm value of isoform A / sum of tpm values of all isoforms * 100).

To examine expression of isoforms in PDAC, we performed western blot analysis of p73 in a panel of PDAC cell lines using a pan-p73 antibody. This revealed the presence of one major band that corresponded to the molecular weight of the longest p73-TA isoform (i.e. p73-TAα) in the majority of the cell lines. Additional lower molecular weight bands in some samples suggested expression of other isoforms, albeit less ubiquitously. Expression levels of p73 isoforms varied across samples, with HS766T and KP2 expressing the highest levels of p73 isoforms (Fig. 1 D). Next, we measured mRNA levels of *TP73-TA* and *TP73-dN* via quantitative reverse transcription PCR (RT-qPCR) using isoform-specific primers. Consistent with the western blot analysis, *TP73-TA* was expressed more highly and ubiquitously compared to *TP73-dN*. One exception to this was HS766T in which *TP73-dN* was expressed at a higher level compared to *TP73-TA* (Supplementary Figure 1 C). We also mined the CCLE and human PDAC organoid transcriptome data sets in which isoform expression data were available. 30/31 cell lines in the CCLE and 39/39 tumor organoids expressed higher levels of *TP73-TA* compared to *TP73-dN*. Specifically, both data sets implicated *TP73-TAα* as the majorly expressed isoform (Fig. 1 E, Supplementary Figure 1 D). Together, these analyses demonstrated that p73-TA is predominantly expressed in PDAC and that it serves as an indicator for disease progression.

### CRISPR-based activation allows moderate upregulation of p73

To examine the functional role of p73 in PDAC, we ectopically expressed p73 by lentiviral delivery of cDNA of *TP73-TAα* in three cell lines—ASPC1, CFPAC1, and PANC1—that express endogenous p73 at low levels (Fig. 2 A). However, we noticed that these cells underwent severe growth arrest compared to the control transduced cells (Fig. 2 B). This phenotype was consistent with previous reports that p73-TA induces p53-like tumor suppressive responses by transactivating classical p53 target genes, such as *CDKN1A* and *GADD45* and inducing cell cycle arrest (Ramadan et al., 2005; Yang & Mckeon, 2000). Such a strong growth arrest phenomena limited us from carrying out downstream experiments. In addition, we found it somewhat paradoxical that *TP73* was widely expressed across different PDAC samples despite its tumor suppressive function. This led us to hypothesize that p73 has additional functions when expressed at moderate levels. To this end, we turned to CRISPR-based activation system (CRISPRa), in which level of gene induction can be carefully controlled by considering different single-guide RNAs (sgRNAs) (Horlbeck et al., 2016). This system has the added advantage of permitting expression of endogenous C-terminus spliced isoforms whose expression patterns slightly vary between samples (Fig. 1 E). We chose sgRNAs targeting upstream of the *TP73* P1 promoter that were able to induce p73-TA at lower levels than those transduced by cDNA, yet to a similar extent as endogenously expressed p73 in KP2 (Fig. 2 A, Supplementary Figure 3 A, B). Notably, these cells slowed down growth to a lesser extent when p73-TA was induced by lentiviral cDNA (Fig. 2 B). In addition, cells stably expressing p73-TA via CRISPRa could be passaged multiple times (>20 passages) in culture (data not shown). To molecularly validate this phenotype, we conducted RNA-sequencing (RNA-seq) experiments on both cDNA and CRISPRa induced samples. We first performed gene ontology analysis on genes that were significantly upregulated in the cDNA induced samples. As expected, we noted that terms related to p53 pathway, apoptosis, and negative regulation of cell proliferation were significantly enriched (Fig. 2 C). Next, we compared the extent of upregulation of the genes that belonged to the aforementioned gene ontologies between cDNA and CRISPRa induced samples. Importantly, cDNA induced samples displayed much stronger upregulation of these genes compared to the CRISPRa induced samples (Fig. 2 D and Supplementary Table 2). Collectively, these data illustrate dose-dependent effect of p73-TA in inducing the anti-proliferative program in PDAC cells. Additionally, these data suggest that CRISPRa is a suitable platform to study the disease-relevant, moderate upregulation of p73-TA seen in PDAC. Accordingly, all subsequent gain-of-function studies were carried out employing CRISPRa.

**Figure 2.**
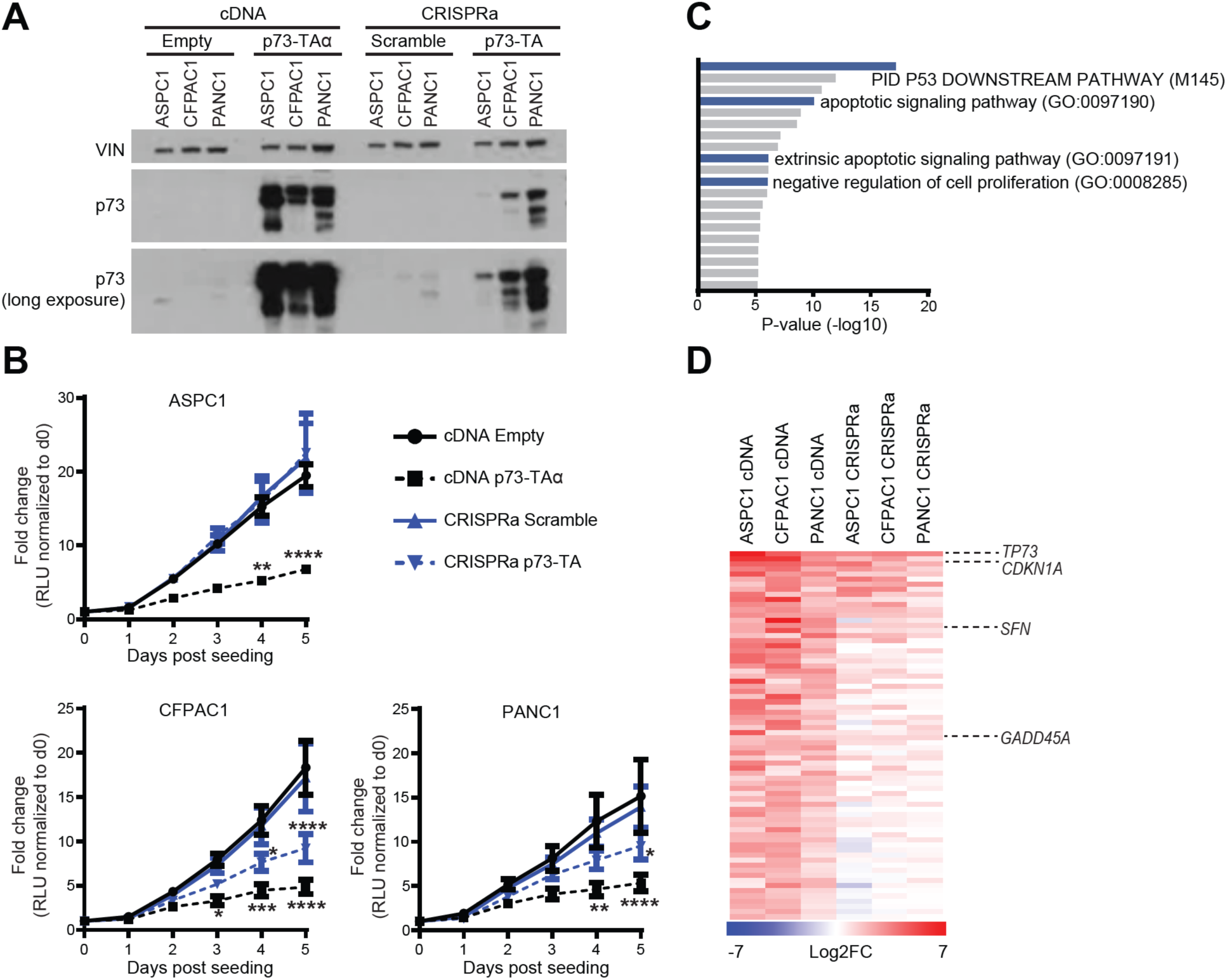
p73-TA elicits p53-like responses in a dose-dependent manner. (*A*) Western blot analysis of p73 following lentiviral delivery of *TP73-TAα* cDNA or CRISPRa-based induction of p73-TA in ASPC1, CFPAC1, and PANC1 cell lines, compared to the empty vector (Empty) or scramble sgRNA (Scramble) controls, respectively. VINCULIN (VIN) shown as a loading control. (*B*) Proliferation of cell lines mentioned in (A) measured by CellTiterGlo Luminescent Cell Viability Assay; luminescence is measured in relative light units (RLU). For each sample, every data point is normalized to the RLU value on day 0 and represented as fold-change. P-values compared to cDNA Empty sample calculated by two-way ANOVA with Sidak’s multiple comparisons test. *, P ≤ 0.05; **, P ≤ 0.01; ***, P ≤ 0.001; ****, P ≤ 0.0001. Mean ± SEM shown (n=3). (*C*-*D*) RNA-seq analysis of cell lines mentioned in (A). (*C*) Gene ontology (GO) analysis using Metascape (Zhou et al., 2019), interrogating significantly upregulated genes following overexpression of p73-TAα using cDNA. Terms are ranked by their p-values; *TP53* and anti-proliferative/apoptotic terms are highlighted in blue. (*D*) Comparison between cDNA and CRISPRa induced samples of 72 genes that were categorized to the gene ontology terms highlighted in (C). Heatmap shows log2 fold-change in the expression levels of genes when p73-TA is overexpressed compared to the controls. *TP73* and example *TP53* target genes are labeled (Zhan et al., 1998; Hermeking et al., 1997; El-Deiry et al., 1993). See also Materials and Methods and Supplementary Table 2.

### p73-TA induces the basal/squamous transcriptional program in PDAC accompanied by changes in the enhancer landscape

As mentioned above, *TP73* is preferentially expressed in the basal-like subtype of PDAC (Fig. 1 C). This encouraged us to examine whether p73 is capable of activating the basal/squamous transcriptional signature in PDAC. To test this, we induced expression of p73-TA in seven PDAC cell lines that express low to moderate levels of endogenous p73 (i.e. ASPC1, BXPC3, CFPAC1, MIAPAC2, PANC1, SUIT2, and SW1990), using two independent sgRNAs (Supplementary Figure 3 B). We then performed RNA-seq analysis on those samples followed by gene set enrichment analysis (GSEA) of a gene set that was preferentially enriched in the basal/squamous lineage PDAC (Somerville et al., 2018). Remarkably, p73-TA strongly upregulated these genes in all cell lines examined (Fig. 3 A, left). In converse experiments, we knocked-out p73-TA in seven cell lines that express moderate to high levels of endogenous p73 (i.e. CAPAN2, HS766T, KP2, PATU8902, PATU8988T, SUIT2, and YAPC). We used CRISPR-Cas9 editing and designed two independent sgRNAs that target genomic regions specific to *TP73-TA* without altering the *TP73-dN* isoform (Supplementary Figure 3 A, C). Here, two cell lines that express p73-TA at the highest levels (i.e. HS766T and KP2) displayed significant downregulation of basal/squamous signature genes (Fig. 3 A, right). To examine whether the remaining p73-dN can regulate the basal/squamous transcriptional program, we knocked-out all isoforms of p73 in the same seven cell lines, using two independent sgRNAs targeting DNA binding domain of *TP73*, and repeated the RNA-seq analysis (Supplementary Figure 3 A, C). All isoform knock-out of p73 did not consistently exacerbate nor alleviate the extent of downregulation of the basal/squamous program compared to the p73-TA-specific knock out (Supplementary Figure 4 A). We additionally validated the knock-out experiments in one tumor-derived PDAC organoid line, hM1a, that expresses high level of *TP73*, and achieved similar results (Supplementary Figure 1 D, Supplementary Figure 4 B) (Tiriac et al., 2018). These results illustrate that p73-TA plays a role in activating the basal/squamous transcriptional program (Fig. 1 D, E and Supplementary Figure 1 C, D).

**Figure 3.**
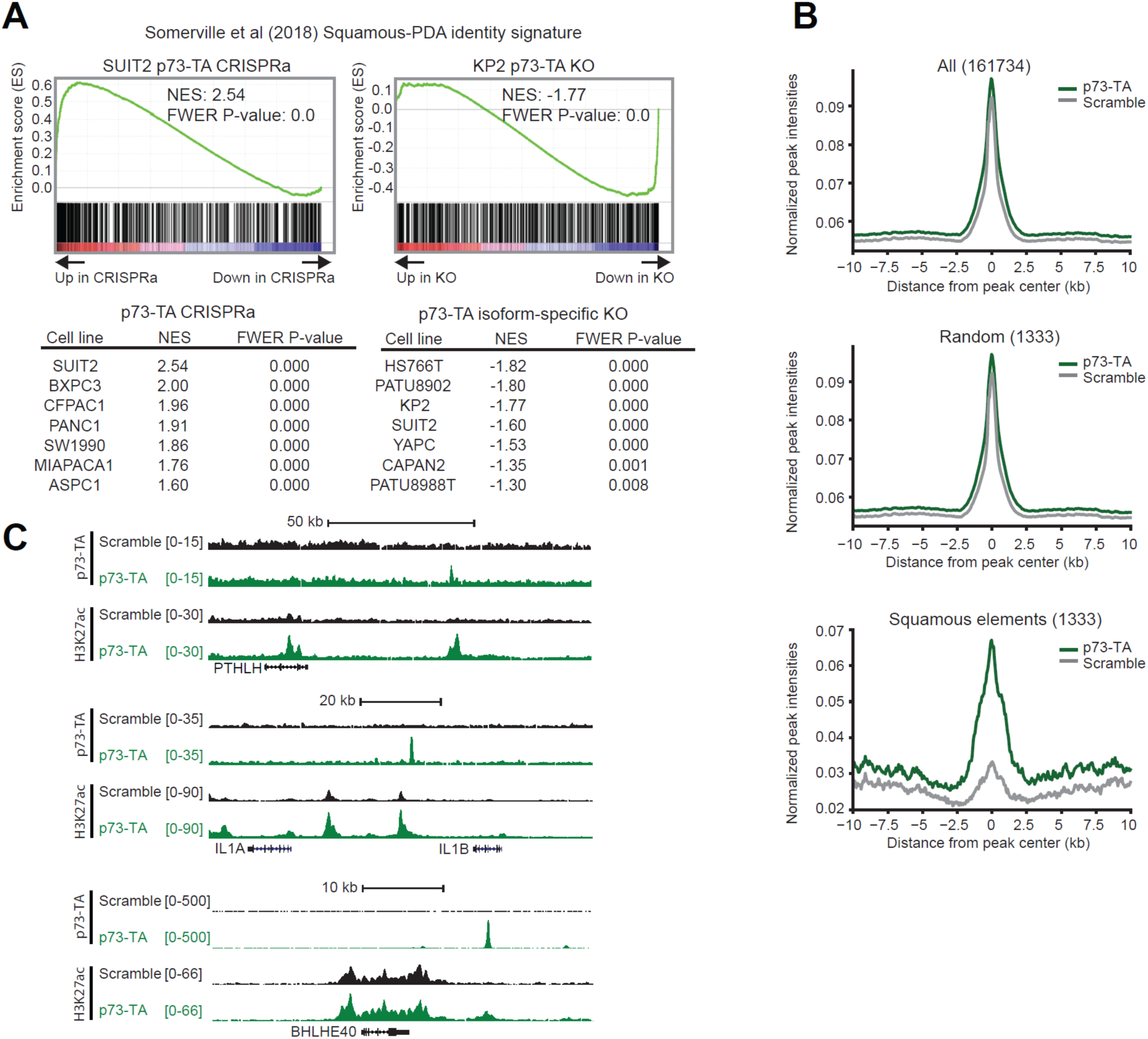
p73-TA promotes squamous transcriptional program in PDAC. (*A*) RNA-seq analysis following activation or ablation of p73-TA in PDAC cell lines. Representative plots and tables summarizing GSEA analysis of the squamous subtype PDAC signature (Somerville et al., 2018) following p73-TA activation (left) or knock-out (right) in a panel of PDAC cell lines. (*B* and *C*) ChIP-seq analysis following p73-TA activation in SUIT2 cell line. (*B*) Metagene plots of H3K27ac signals in all elements (top), random elements (middle), and the squamous elements (bottom) (Somerville et al., 2018). (*C*) ChIP-seq profiles of H3K27ac and p73-TA signals at example genes associated with squamous lineage PDAC and disease aggression (Biffi et al., 2019; Hamdan & Johnsen, 2018; Pitarresi et al., 2021; Somerville et al., 2018; Somerville, Biffi, et al., 2020).

Next, using chromatin immunoprecipitation sequencing (ChIP-seq), we performed analysis of H3K27ac, a histone modification mark enriched in active enhancer elements (Creyghton et al., 2010; Rada-iglesias et al., 2011), following induction of p73-TA in SUIT2 cells. We specifically interrogated the “squamous elements,” genomic regions enriched with H3K27ac preferentially in p63-expressing, basal/squamous lineage PDAC cell lines (Somerville et al., 2018). Complementing our RNA-seq data, we saw a significant and selective increase in H3K27ac at the squamous elements upon induction of p73-TA, while no changes were observed at control regions (Fig. 3 B). To further probe whether p73 directly binds to enhancer elements to drive the basal/squamous program, we performed ChIP-seq analysis of p73, using a p73-TA-specific antibody. Here, we observed p73 binding peaks near genes such as *PTHLH*, *IL1A*, and *BHLHE40*, that are associated with the squamous lineage and disease aggressiveness in PDAC (Fig. 3 C) (Biffi et al., 2019; Hamdan & Johnsen, 2018; Pitarresi et al., 2021; Somerville et al., 2018; Somerville, Biffi, et al., 2020). Together, these results illustrate that p73-TA can activate the basal/squamous program and reprogram enhancers associated with disease progression.

### p73-TA activates a ciliogenesis transcriptional signatures of the ‘Basal B’ subtype of PDAC

A recent study by Chan-Seng-Yue and colleagues (Chan-Seng-Yue et al., 2020) analyzed tumors from the COMPASS trial and reported that the previously designated basal/squamous lineage tumors can be further classified into ‘Basal A’ or ‘Basal B’ subtypes (Chan-Seng-Yue et al., 2020). In this study, excluding the ‘Hybrid’ tumors that could not be classified into a definitive subtype due to the presence of multiple expression signatures, ∼14% of tumors were classified as Basal A and ∼13% as Basal B (Chan-Seng-Yue et al., 2020). Two sets of genes, namely Signature 2 and Signature 10, were significantly expressed in the Basal subtypes; Signature 2 included classical basal/squamous lineage markers such as *TP63*, *KRT5*, and *KRT6A* (Chan-Seng-Yue et al., 2020; Kaufmann et al., 2001; Somerville et al., 2018), and Signature 10 consisted of genes related to ciliogenesis program such as *FOXJ1*, *DRC1*, and *CFAP54* (Chan-Seng-Yue et al., 2020; Marshall et al., 2016; Mckenzie et al., 2015; Nemajerova et al., 2016). While Basal A subtype preferentially expressed Signature 2 genes, Basal B subtype expressed moderate levels of the Signature 2 genes and high levels of the Signature 10 genes. Interestingly, Basal B subtype was classified as less aggressive than the Basal A subtype, proposing that the expression of Signature 10 genes serves as a marker for the less aggressive basal/squamous lineage of PDAC (Chan-Seng-Yue et al., 2020). Previous studies have highlighted a central role of p73 as an activator of FOXJ1—a master regulator of ciliogenesis—and the downstream transcriptional program required for multiciliated cell differentiation in mouse lung epithelium (Brody et al., 2000; Chan-Seng-Yue et al., 2020; Marshall et al., 2016; Nemajerova et al., 2016; Yu et al., 2008). This coincidental observation that a well-described p73-driven transcriptional program was upregulated in the Basal B subtype of PDAC prompted us to examine whether p73 has a causal role in regulating the Signature 10 genes. To do this, we performed GSEA evaluating Signature 2 and Signature 10 in the aforementioned RNA-seq data following p73-TA activation and knock-out. First, we noticed that Signature 2 analysis yielded a very similar result as the Somerville et al. squamous signature analysis (Supplementary Figure 4 C). This result was expected given that ∼21% of genes overlap between the two gene sets (Supplementary Figure 4 D) (Chan-Seng-Yue et al., 2020; Somerville et al., 2018). On the other hand, analysis of Signature 10 led to interesting results. While activation of p73-TA variably affected the expression of the Signature 10 genes across cell lines, knock-out of p73-TA strongly downregulated Signature 10 genes in five out of seven cell lines (Supplementary Figure 4 C). This result suggested that p73-TA is required to maintain the expression of Signature 10 genes in PDAC. Of note, *FOXJ1* is one of the most highly expressed transcription factors in the *TP73*^high^ compared to the *TP73*^low^ groups (Supplementary Figure 4 E), and tumors expressing high levels of *TP73* also expressed *FOXJ1* at high levels (Supplementary Figure 4 F). In addition, knocking out p73-TA abrogated the protein expression of *FOXJ1* (Supplementary Figure 4 G). Together, this analysis demonstrated that 1) the previously described p73-FOXJ1 axis is present in a subset of PDAC samples 2) p73-TA not only promotes the basal/squamous program but also is necessary to maintain the transcriptional network unique to the newly described Basal B subtype of PDAC, which is enriched for high expression of ciliogenesis genes.

### p73-TA promotes PDAC cell migration and invasion

We performed gene ontology analysis of significantly upregulated genes following p73-TA activation and identified cellular processes related to cell migration and invasion, such as regulation of cell adhesion, positive regulation of locomotion, chemotaxis, and actin filament-based process (Fig. 4 A). This result was consistent with previous reports illustrating that p73-TA regulates cell adhesion and migration of various cell types (Sablina et al., 2003; Santos Guasch et al., 2018; Xie et al., 2018). To validate these transcriptional changes, we performed scratch assays on PDAC cell lines subsequent to activation of p73-TA and observed cell line-dependent responses in wound closure. p73-TA accelerated wound closure most strongly in SUIT2 and SW1990 (Fig. 4 B), and moderately but significantly in MIAPACA2 and PANC1 (Supplementary Figure 5 A). No effect was observed in ASPC1, BXPC3 and CFPAC1 (data not shown). We next conducted transwell assays in MIAPACA2, PANC1, SUIT2, and SW1990 to measure invasive ability of cells through extra cellular matrix. In MIAPACA2, PANC1, and SW1990, p73-TA promoted invasion of cells, as indicated by increase in percent cells that invaded through matrigel-coated membranes over cells that freely migrated through uncoated membranes (Fig. 4 C). There was no significant change in percent invasion of SUIT2 cells, but this was mainly driven by robust increase in migratory ability of cells when p73-TA was activated, regardless of the presence of the extra cellular matrix (Fig. 4 C). Lastly, p73-TA activated cells plated in semisolid media to encourage anchorage-independent growth formed more invasive looking colonies compared to the control cells (Fig. 4 D). Overall, these results demonstrated the ability of p73-TA to promote cell migration and invasion in a subset of PDAC cell lines.

**Figure 4.**
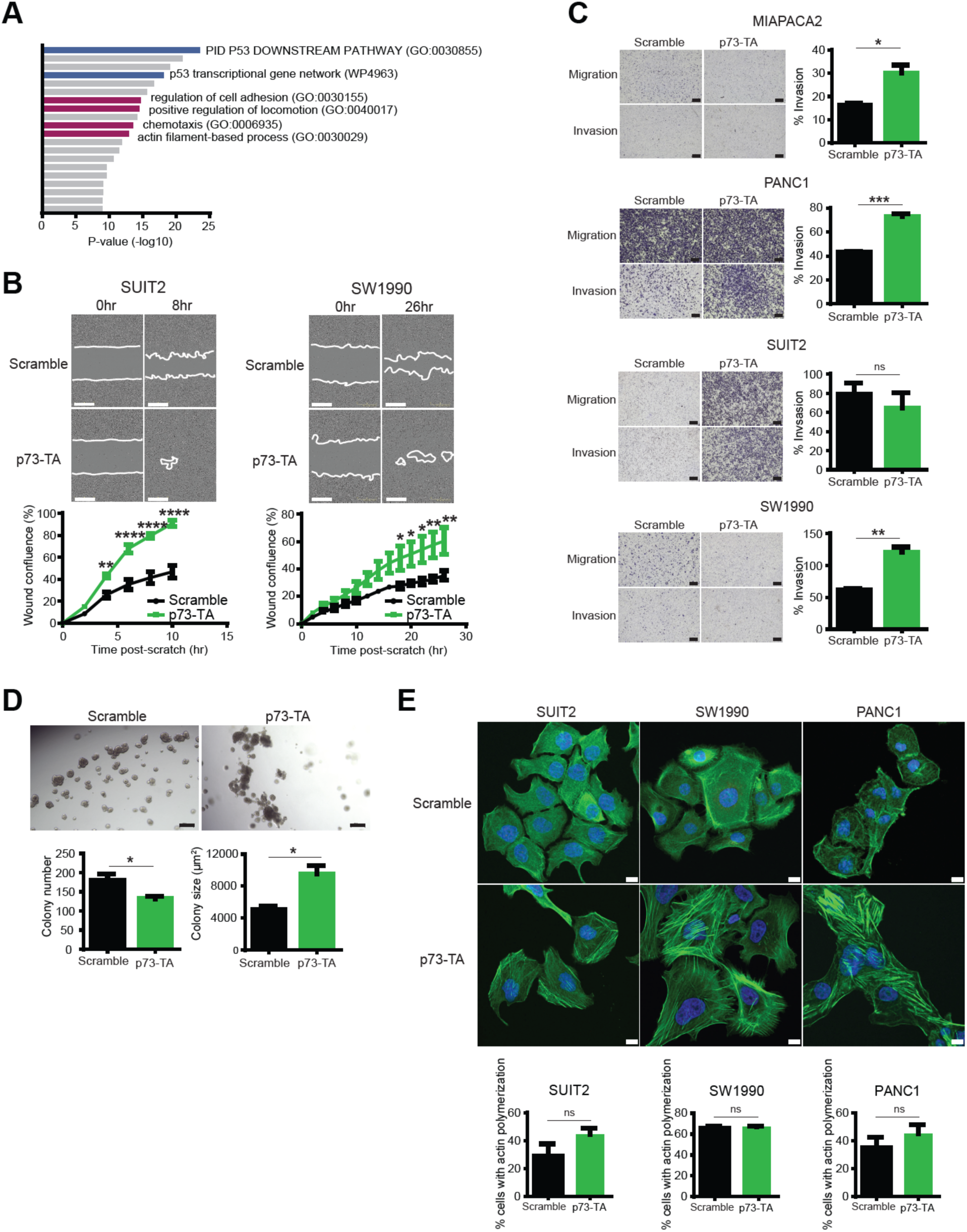
p73-TA promotes cell migration and invasion in PDAC cells. (*A*) GO analysis using Metascape (Zhou et al., 2019), interrogating significantly upregulated genes following activation of p73-TA in a panel of PDAC cell lines. Terms are ranked by their p-values; *TP53*-related terms are highlighted in blue and cell invasion/migration-related terms are highlighted in magenta. (*B*) Scratch assays following activation of p73-TA compared to the control (Scramble). Quantification of % wound confluence plotted at the indicated time points post-scratch and representative images are shown; scale bar indicates 400 µm. P-values compared to Scramble calculated using two-way ANOVA with Sidak’s multiple comparisons test. Mean ± SEM shown (n=3). *, P ≤ 0.05; **, P ≤ 0.01; ****, P ≤ 0.0001. (*C*) Transwell assays following activation of p73-TA compared to the control (Scramble). Bar charts show % invasion calculated by number of cells migrated through Matrigel-coated membrane (Invasion) vs. uncoated membrane (Migration). Representative images shown; scale bar indicates 200 µm. P-values calculated using unpaired t-test. Mean ± SEM shown (n=3). ns, P > 0.05; *, P ≤ 0.05; **, P ≤ 0.01; ***, P ≤ 0.001. (*D*) 3D Matrigel colony formation assays following activation of p73-TA in SUIT2 cells compared to the control (Scramble). Bar charts show colony number and size. P-values calculated using unpaired t-test. Mean ± SEM shown (n=3). *, P ≤ 0.05. Representative images shown; scale bar indicates 200 µm. (*E*) Representative images of phalloidin staining (green) to visualize F-actin following p73-TA activation compared to the control (Scramble); scale bar indicates 10 µm. Nuclei stained with DAPI (blue). See also Supplementary Figure 6. Bar charts show quantification of % cells presenting actin polymerization. P-values calculated using unpaired t-test. Mean ± SEM shown (n=3). ns, P > 0.05.

Actin cytoskeleton reorganization is closely associated with cell migration (Tojkander et al., 2012; Vallenius, 2013). Assembly and dynamics of actin filaments regulate contractile force, interaction with focal adhesions, and mechanosensing of migrating cells (Parsons et al., 2010; Prager-khoutorsky et al., 2011; Tojkander et al., 2012). To visualize actin filaments in cell lines that displayed strong migratory and/or invasive phenotype, we performed phalloidin staining in PANC1, SUIT2, and SW1990 cells, following p73-TA activation (Fig. 4 E, Supplementary Figure 6). While there was no significant change in the proportion of cells that presented actin polymerization, p73-TA-activated cells formed actin filament bundles that appeared more orderly and dynamic compared to the control cells. Notably, these actin bundles were predominantly formed at the leading edge of p73-TA-activated SUIT2 and SW1990 cells (Fig. 4 E, Supplementary Figure 6). Consistently, development of such a morphological asymmetry (i.e. cell polarization) is one of the basic principles followed by migrating cells (Gómez-moutón, 2007; Vallenius, 2013).

Rho GTPases and integrins are well-appreciated regulators of cell migration and invasion and are often taken advantaged by cancer cells to promote disease progression (Guo & Giancotti, 2004; Hamidi & Ivaska, 2018; Heasman & Ridley, 2008; Lawson & Ridley, 2018). Building on the *in vitro* migration/invasion/actin remodeling phenotypes induced by p73-TA, we mined our RNA-seq data to see whether p73-TA can regulate the expression of Rho GTPases and/or integrins. Remarkably, a subset of Rho GTPases and integrins were upregulated upon p73-TA activation across cell lines (Supplementary Figure 5 B). We also validated in patient transcriptome data that some of these genes such as *RND3*, *RHOD*, *RAC2*, *ITGA3*, *ITGA6*, and *ITGB4* were more highly expressed in the *TP73*^high^, compared to the *TP73*^low^ tumors (Supplementary Figure 5 C). Though function of these genes in regulating PDAC cell migration and invasion will need to be experimentally tested, our analysis demonstrated that p73-TA can broadly promote cell migration and invasion and the expression of Rho GTPases and integrins in PDAC cells.

### p73-TA confers dependency in a subset of PDAC cell lines

We noted that two cell lines that express highest levels of p73 (i.e. KP2 and HS766T) are dependent on *TP73* for their growth as demonstrated in the DepMap data (Fig. 5 A) (Meyers et al 2017 and Dempster et al 2019). We validated this result by performing GFP competition assays, using two independent sgRNAs that target all isoforms of p73 and two independent sgRNAs that specifically target p73-TA (Fig. 5 B). Notably, we observed that knocking-out p73-TA is sufficient to confer growth disadvantage to these cells, highlighting the essential role of p73-TA in regulating the *TP73*-addiction of PDAC cells.

**Figure 5.**
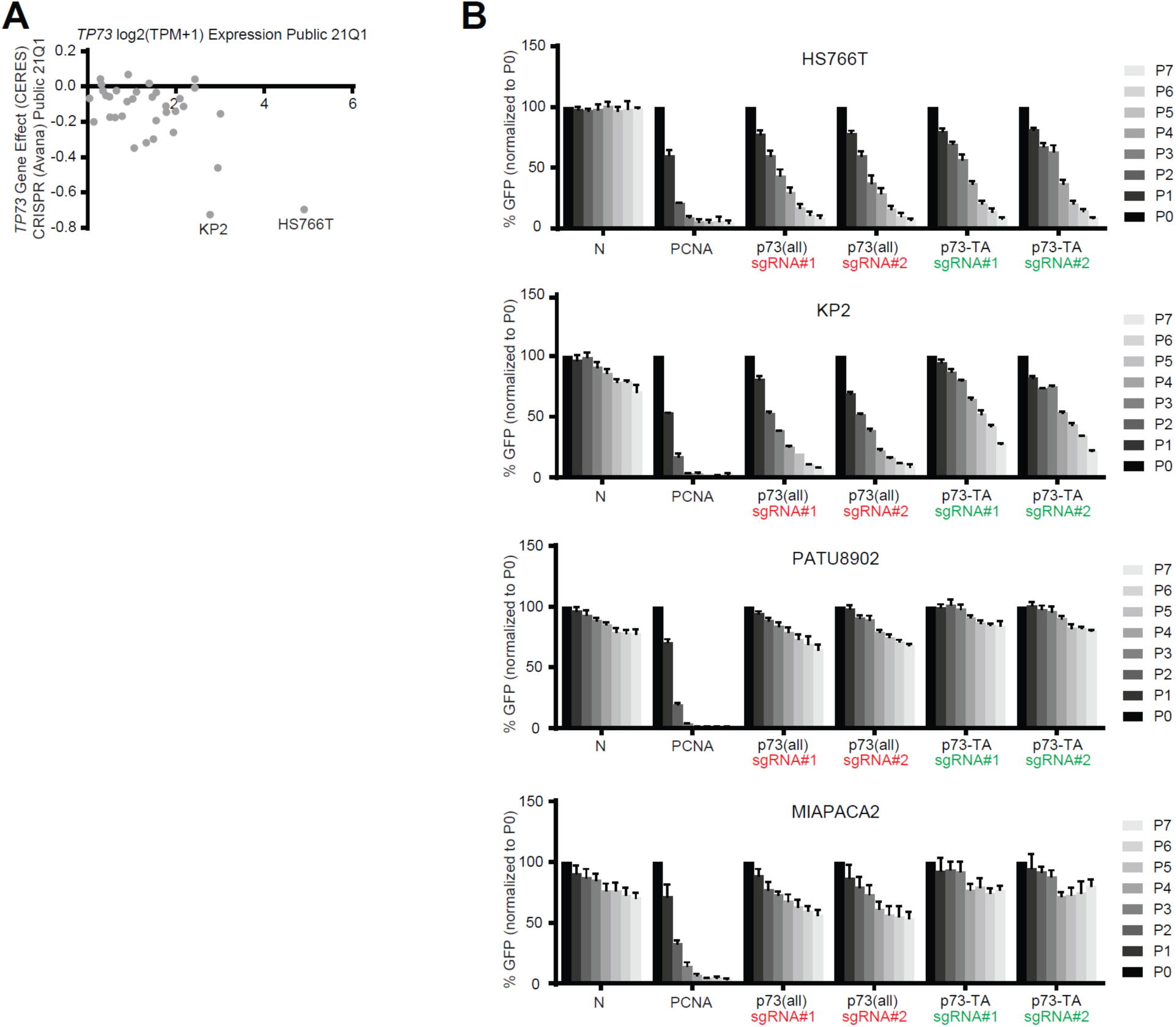
A subset of PDAC cells depend on p73-TA for growth. (*A*) Illustration of DepMap data (Meyers et al., 2017 and Dempster et al., 2019) focusing on expression and dependency score (CERES) of *TP73* in 34 PDAC cell lines. CERES score less than -0.5 considered as dependent, and the two dependent cell lines, KP2 and HS766T, are labeled. (*B*) Validation of *TP73* dependency using competition-based cell proliferation assays in *TP73*-dependent (KP2 and HS766T) and *TP73*-neutral (PATU8902 and MIAPACA2) cell lines. Cas9-expressing cell lines were infected with sgRNAs whose expression plasmids also encode GFP. Non-targeting sgRNAs (N), sgRNA targeting *PCNA* (positive control), sgRNA targeting all isoforms of p73 (sgRNA #1 and #2, red), and sgRNAs targeting p73-TA (sgRNA #1 and #2, green) were used. Percentage of GFP+ cells was measured every 2 or 3 days (P1-P7) starting day 3 (P0) post-sgRNA infection. For each sgRNA experiment, every data point is normalized to the percent GFP value at P0. See also Supplementary Figure 3 A.

## Discussion

In this study, we provide evidence supporting that transcription factor p73-TA plays oncogenic roles in PDAC. Despite previous reports illustrating that p73-TA is generally tumor suppressive by activating p53 target genes (Melino et al., 2002; Zhu et al., 1998), there are clues suggesting that p73-TA is not a classical tumor suppressor, at least in the context of PDAC: *TP73* is rarely mutated in cancers including PDAC (Supplementary Figure 6 B) (Dötsch et al., 2010); *TP73-TA* is widely detected across PDAC tumors, cell lines, and organoids; and high expression of *TP73* is correlated with worse clinical outcome. Thus, we hypothesize that PDAC tumors activate expression of p73-TA to promote disease progression.

Dose-dependent regulation of transcription factors during lineage commitment has been demonstrated in both development and disease (Doghman et al., 2013; Niwa et al., 2000; Simmons et al., 2012; Takeuchi et al., 2005). As a classical example, precise level of Oct3/4 is necessary to main pluripotency of mouse embryonic stem cells, while reduced or elevated level of the protein leads to aberrant lineage commitment (Niwa et al., 2000). In another study, high or low expression of Pax5 in early lymphoid progenitors leads to normal B-lineage development or leukemia, respectively (Simmons et al., 2012). We demonstrate that the tumor suppressive effect of p73-TA is dose-dependent in PDAC cells. When p73-TA is overexpressed to supraphysiological level, cells experience severe growth retardation. On the contrary, cells expressing disease-relevant level of p73-TA moderately activate p53-like responses but continue to proliferate; these cells also take on a basal/squamous lineage program associated with disease aggression. This observation reemphasizes the significance of dosage as well as pleiotropic ability of transcription factors during normal and malignant lineage reprogramming.

*TP73* is expressed at a higher level in the basal-like/squamous subtype of PDAC compared to the progenitor/classical subtype. Consistent with this observation, we show that p73-TA can upregulate the basal/squamous program in PDAC cells by remodeling the enhancer landscape. This adds to the growing evidence that chromatin remodeling is a critical aspect of lineage reprogramming in PDAC (Andricovich et al., 2018; Roe et al., 2017; Somerville et al., 2018; Somerville, Xu, et al., 2020). In addition, p73-TA knock-out leads to a significant downregulation of the basal/squamous program in two cell lines that express p73-TA at high levels, suggesting that p73-TA maintains the basal/squamous lineage program in a subset of PDAC cells. We also note that the newly described Basal B signature of PDAC, enriched with genes related to ciliogenesis (Chan-Seng-Yue et al., 2020), is specifically downregulated by knocking-out p73-TA across PDAC cell lines. This observation is highly consistent with the findings from Nemajerova et al. and Marshall et al., that p73-TA is a central regulator of multiciliated cell differentiation (Marshall et al., 2016; Nemajerova et al., 2016). However, the role of ciliogenesis and/or the function of the Basal B signature genes in PDAC biology remain to be determined.

Given that the p63 isoform that lacks the N-terminus transactivation domain (p63-dN) is a well-described master regulator of the basal/squamous lineage PDAC (Andricovich et al., 2018; Hamdan & Johnsen, 2018; Somerville et al., 2018) and that it belongs to the same paralog family as p73, the important question remains: what is the relationship, if any, between p73-TA and p63-dN in regulating the basal/squamous lineage in PDAC. Addressing this question will first require single-cell level examination of expression of each factor in PDAC cells. Of note, a study by Marshall et al. provides interesting example of cooperation between p73 and p63 during lung epithelial development (Marshall et al., 2016). Here, the authors demonstrate that perturbations of p73 or p63 leads to distinct changes in the composition of cell types in lung epithelium. For example, p73 knock-out not only results in reduction of ciliated cells but also basal cells; p63 knock-out results in reduction of the basal cells but increase in ciliated cells. One possible explanation for this phenotype is that a subset of basal cells is maintained by p73, and/or that p63 represses ciliated cell development (Marshall et al., 2016). Given that p73 and p63 bind and regulate overlapping set of genes (Harms et al., 2004; Marshall et al., 2016), the authors propose that changes in the stoichiometry of the two proteins and/or their isoforms may be one mechanism through which p63 and p73 together determine the identity of lung epithelium (Marshall et al., 2016). Based on the presence of both p63-regulated basal/squamous lineage program and p73-regulated basal/squamous/ciliogenesis program in PDAC and lung epithelial cells, it is tempting to speculate that a similar co-regulatory mechanism of p73 and p63 exists in the basal/squamous lineage PDAC. Perhaps, the Basal A and Basal B subtypes identified by Chan-Seng-Yue et al. demonstrate manifestation of such a regulation (Chan-Seng-Yue et al., 2020).

We show that p73-TA can regulate cell migration and invasion, which is associated with reorganization of actin cytoskeleton in PDAC cells. This is consistent with previous reports that p73-TA can promote cell migration and invasion (Landré et al., 2016; Sablina et al., 2003). We also demonstrate that p73-TA can regulate expression of Rho GTPases and integrins, which are well-known regulators of tumor invasion and metastasis (Guo & Giancotti, 2004; Jung et al., 2020; Lawson & Ridley, 2018). For instance, integrin α6β4 is broadly overexpressed in PDAC and promotes disease aggression (Carpenter et al., 2015; Cruz-Monserrate & O’Connor, 2008). In our study, we find that p73-TA upregulates expression of both *ITGA6* and *ITGB4*, and that both genes are more highly expressed in the *TP73*^high^ tumors. Given that regulation and function of Rho GTPases and integrins in cancers are complex and are context-dependent (Guo & Giancotti, 2004; Jung et al., 2020; Lawson & Ridley, 2018), function of individual p73-TA-regulated Rho GTPases and integrins in PDAC will need to be empirically validated. Nevertheless, our observations point out various ways p73-TA can promote cell migration and invasion of PDAC cells.

In summary, this study reports dose-dependent oncogenic roles of p73-TA in PDAC. These findings are in support of a view that p73-TA does not act as a classical tumor suppressor like p53 and an observation that *TP53* mutations are common in cancers while mutations of *TP73* are rare. However, our findings also not support that p73-TA is a a powerful oncogene due to its inherent ability to activate p53 target genes. In fact, we speculate that p73-TA may act in a dichotomous manner in PDAC cells with a scenario in which moderate expression of p73-TA confers and maintains the basal/squamous lineage program of PDAC cells, promotes cell migration and invasion, but attenuates proliferative ability. However, perplexing questions still remains as to why PDAC cells become addicted to p73-TA, bypassing excessive activation of the p53 target genes. To this end, examining the mechanism of addiction to p73 and/or upstream signal leading to upregulation of p73 in PDAC cells will provide further insights into the role of this lineage master regulator in PDAC biology.

## Authors contributions

S.K. Hur: Conceived the project, designed and performed the experiments, and wrote the manuscript. T.D.D. Somerville: Assisted in TP73 dependency validation experiments. X.S Wu: Analyzed ChIP-seq data. D. Maia-Silva: Provided reagents for CRISPRa studies. O.E. Demerdash: Analyzed PDAC organoid TP73 isoform expression data. D.A. Tuveson: Provided PDAC organoid transcriptome data and shared hM1a PDAC organoid. F. Notta: Shared the COMPASS trial data (Chan-Seng-Yue et al., 2020) and provided feedback on the Basal B subtype characteristics. C.R. Vakoc: Supervised the study, acquired fundings, and wrote the manuscript.

## Acknowledgements

This work was supported by Cold Spring Harbor Laboratory NCI Cancer Center Support grant CA045508. Additional funding was provided to C.R.V. by the Pershing Square Sohn Cancer Research Alliance, National Institutes of Health grants CA013106 and CA229699, and the Cold Spring Harbor Laboratory and Northwell Health Affiliation. D.M.-S. was supported by a Boehringer Ingelheim Fonds Ph.D. fellowship.

---

**Supplementary Figure 1.**
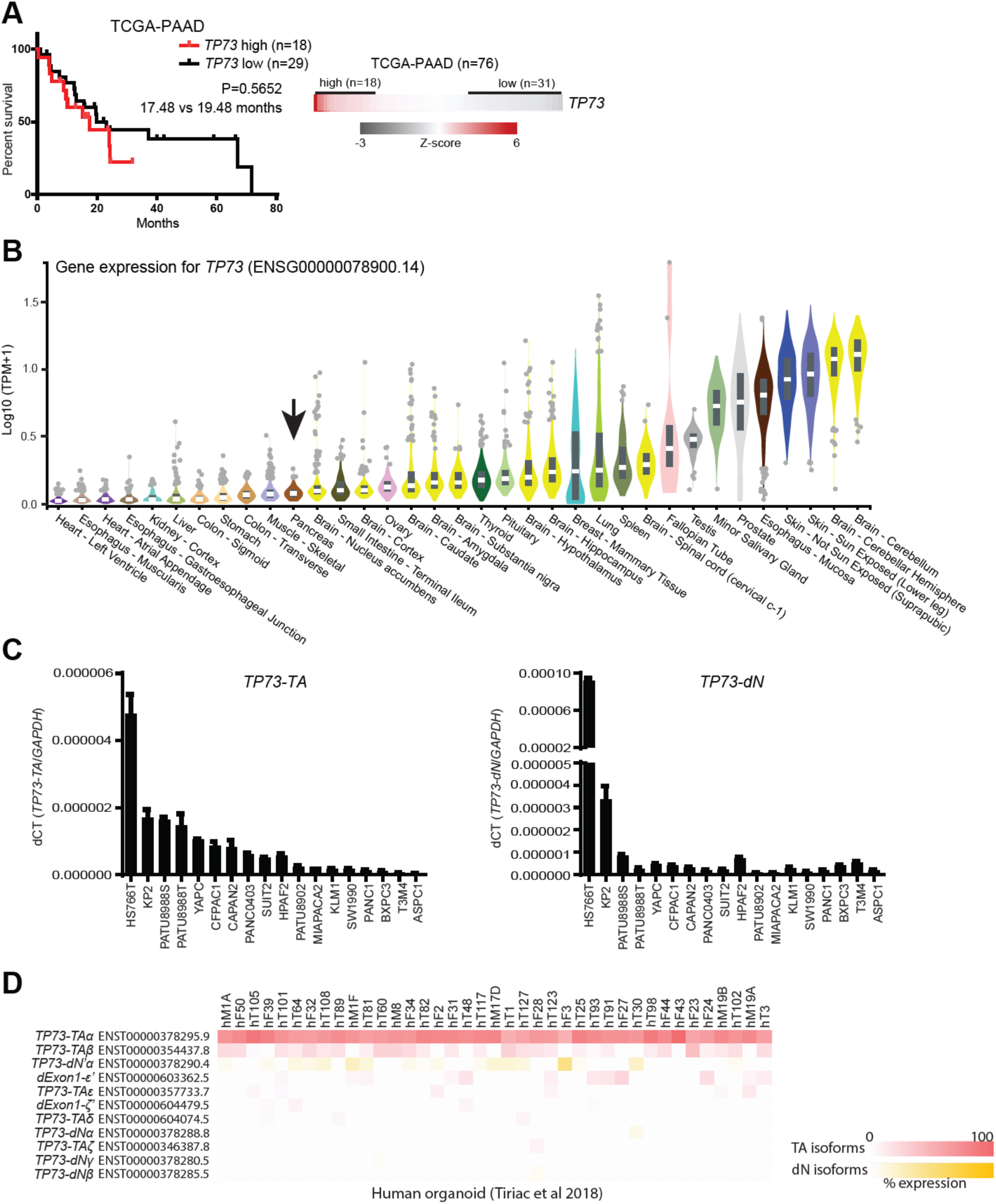
*TP73* is prognostic and expresses multiple isoforms; *TP73-TA* is predominantly expressed in PDAC. Related to Figure 1. (*A*) Survival curve of patients stratified by *TP73* expression. Log-rank (Mantel-Cox) test used to assess the median survival values and p-values. (*B*) Expression of *TP73* in normal tissues from the GTEx project (The GTEx Consortium, 2015). Arrow points to pancreas tissue. (*C*) Expression of *TP73* isoforms in PDAC cell lines analyzed by RT-qPCR using *TP73-TA* specific primers (left) or *TP73-dN*-specific primers (right). Expression levels of *TP73* isoforms were measured relative to the expression level of *GAPDH*, and presented as dCT (i.e. 2 ^ -dCT) values. Mean ± SEM shown (n=3). (*D*) *TP73* isoform expression in PDAC organoids derived from primary tumors (hT), metastases (hM), or fine needle biopsies of primary or metastatic lesions (hF) (Tiriac et al., 2018). For the heatmap, organoids are listed in descending order (from left to right) of total *TP73* isoform expression. See figure legend for Figure. 1 E for additional details.

**Supplementary Figure 2.**
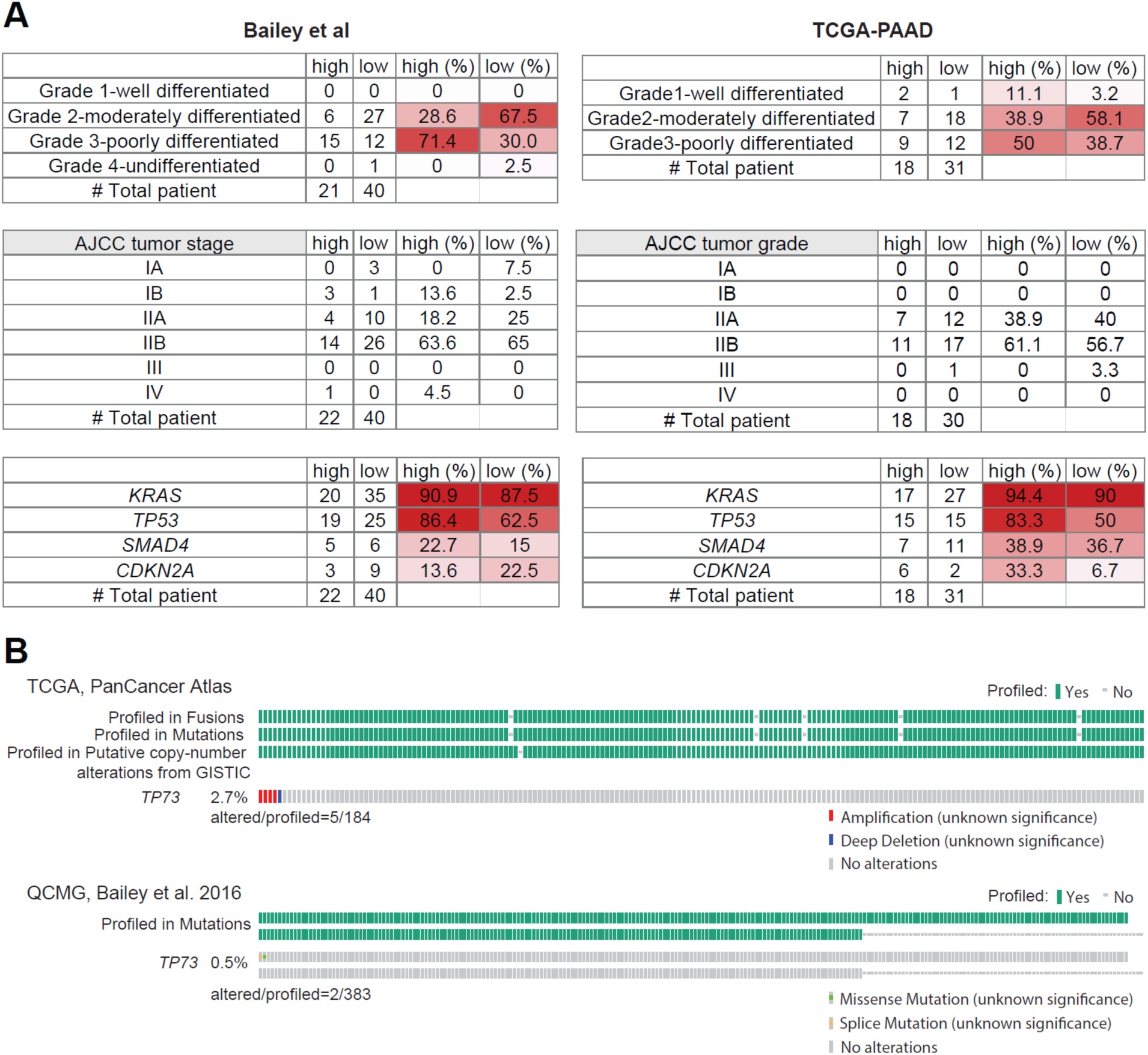
*TP73*^high^ tumors are poorly differentiated and present more frequent *TP53* mutations compared to the *TP73*^low^ tumors, while *TP73* is rarely mutated in PDAC. Related to Figure 1. (*A*) Comparison between *TP73*^high^ (high) and *TP73*^low^ (low) tumors from the indicated studies, regarding tumor histology, tumor stage/grade, and commonly mutated genes in PDAC. (*B*) *TP73* mutational analysis from the indicated studies downloaded from the cBioPortal (Cerami et al., 2012).

**Supplementary Figure 3.**
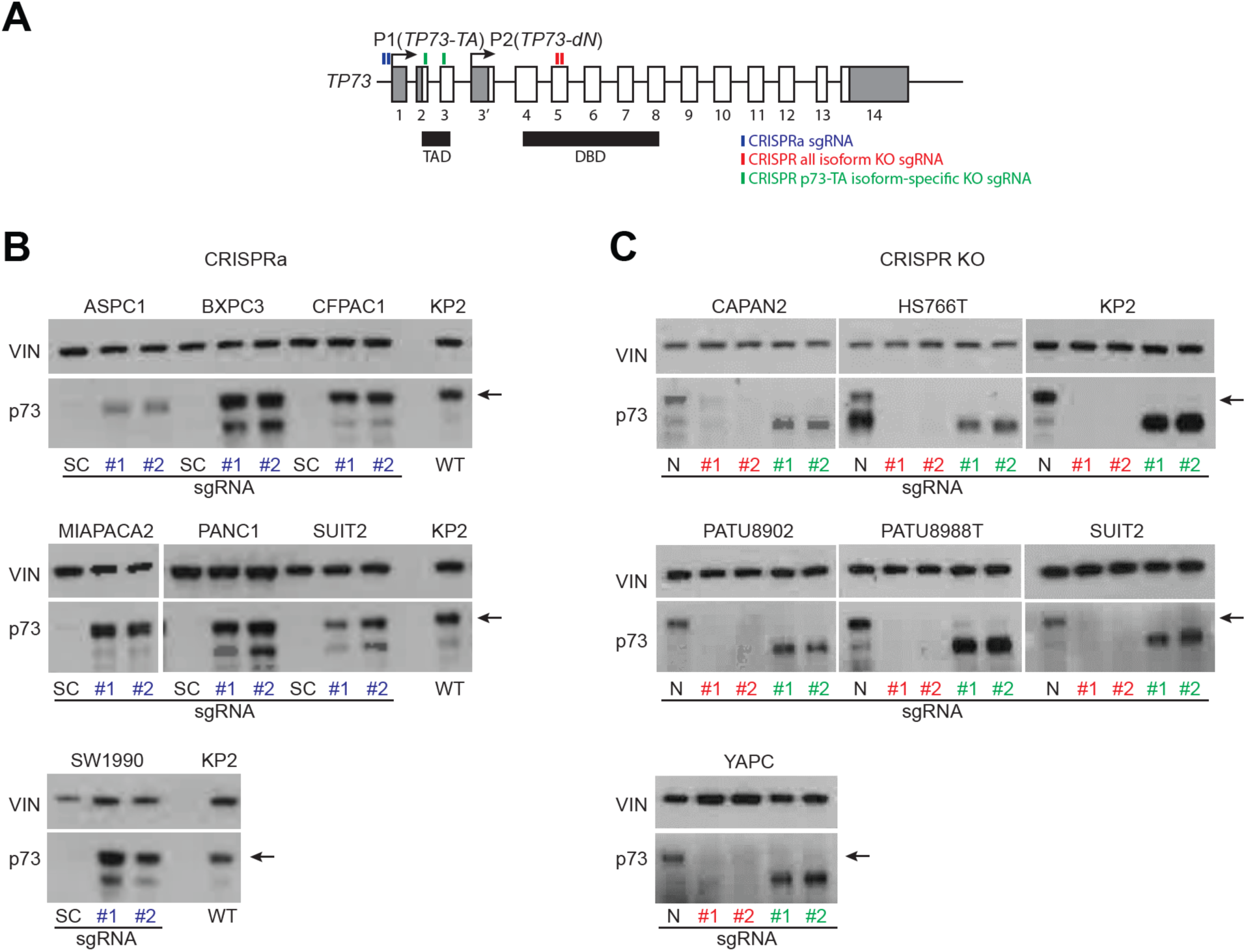
CRISPR-based activation and knock-out of p73. Related to Figure 2 and 3. (*A*) Schematic of the human *TP73* gene and regions targeted by sgRNAs used in this study. Exons (white boxes), UTRs (gray boxes), and introns (thin lines) are depicted. Exons are numbered. Transactivation domain (TAD) and dna-binding domain (DBD) are indicated with black boxes. sgRNAs used for CRISPRa-based induction of p73-TA (blue), CRISPR-based knock-out of all isoforms of p73 (red), CRISPR-based knock-out of p73-TA (green) are illustrated. (*B* and *C*) Western blot analysis of p73 using a pan-p73 antibody following activation or ablation p73 in a panel of PDAC cell lines. Bands corresponding to the molecular weight of the longest p73 isoform (i.e. p73-TAα) indicated with arrows. VINCULIN (VIN) shown as a loading control. (*B*) CRISPRa-based induction of p73-TA using a scramble control (SC) or two independent sgRNAs targeting upstream of P1 promoter (#1 and #2, blue). Wild-type (WT) KP2 cell line included as a reference for endogenous p73 expression. (*C*) CRISPR-based knock-out of p73 using a non-targeting sgRNA (N), two independent sgRNAs targeting DBD (#1 and #2, red) or two independent sgRNAs targeting TAD (#1 and #2, green).

**Supplementary Figure 4.**
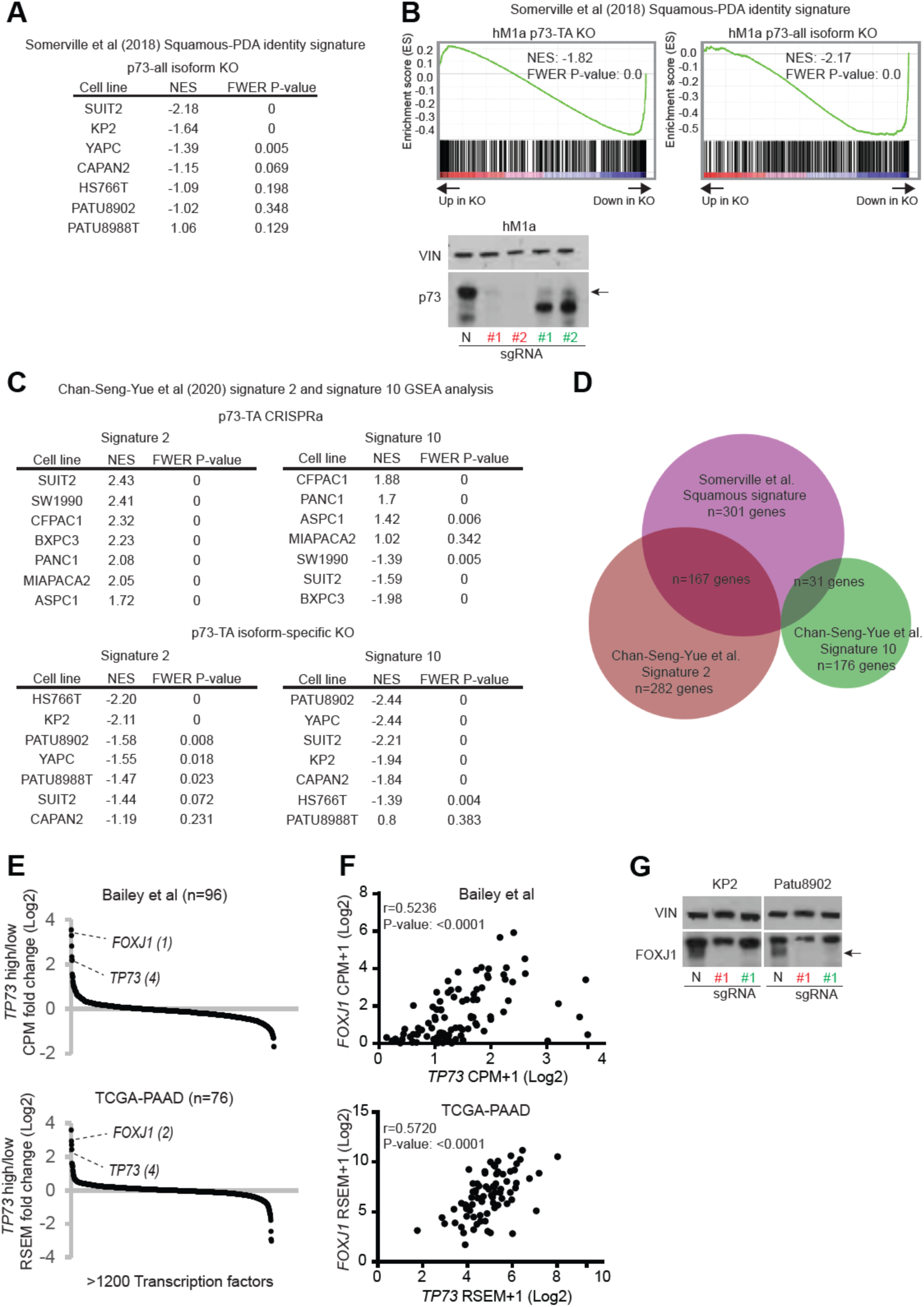
p73-TA promotes squamous program and maintains transcriptional program associated with a newly described ‘Basal B’ subtype of PDAC. Related to Figure 3. (*A*) RNA-seq analysis following ablation of p73 in PDAC cell lines. Table summarizing GSEA analysis of the squamous subtype PDAC signature (Somerville et al., 2018) subsequent to knocking-out all isoforms of p73. (*B*) Western blot and RNA-seq analysis following p73 knock-out in hM1a PDAC organoid. GSEA plot evaluating the squamous subtype PDAC signature (Somerville et al., 2018) after knocking-out p73-TA (left) or all isoforms of p73 (right). For western blot, sgRNAs targeting all isoforms of p73 (#1and #2, red), sgRNAs targeting p73-TA (#1 and #2, green), and non-targeting sgRNA (N) are shown. Bands corresponding to the molecular weight of the longest p73 isoform (i.e. p73-TAα) indicated with an arrow. VINCULIN (VIN) shown as a loading control. (*C*) Tables summarizing GSEA analysis of signature 2 and signature 10 (Chan-Seng-Yue et al., 2020) after activating (top) or knocking-out (bottom) p73-TA in a panel of PDAC cell lines. (*D*) Venn diagram showing number of genes overlapping between Somerville et al. squamous signature (Somerville et al., 2018), and signature 2 and signature 10 from Chan-Seng-Yue et al. (Chan-Seng-Yue et al., 2020). (*E*) Expression of transcription factors in *TP73*^high^ vs *TP73*^low^ tumors. Transcription factors are ranked by their mean log2 fold-change in expression levels in *TP73*^high^ vs. *TP73*^low^ tumors. Rank of each gene is written inside parentheses. (*F*) Pearson correlation analysis of expression levels between *TP73* and *FOXJ1*. Pearson correlation coefficient (r) and P-value shown. (*G*) Western blot analysis of FOXJ1 subsequent to knocking-out p73. sgRNA targeting all isoforms of p73 (#1, red), sgRNA targeting p73-TA (#1, green), and non-targeting sgRNA (N) are shown. Bands corresponding to the molecular weight of FOXJ1 indicated with an arrow. VINCULIN (VIN) shown as a loading control.

**Supplementary Figure 5.**
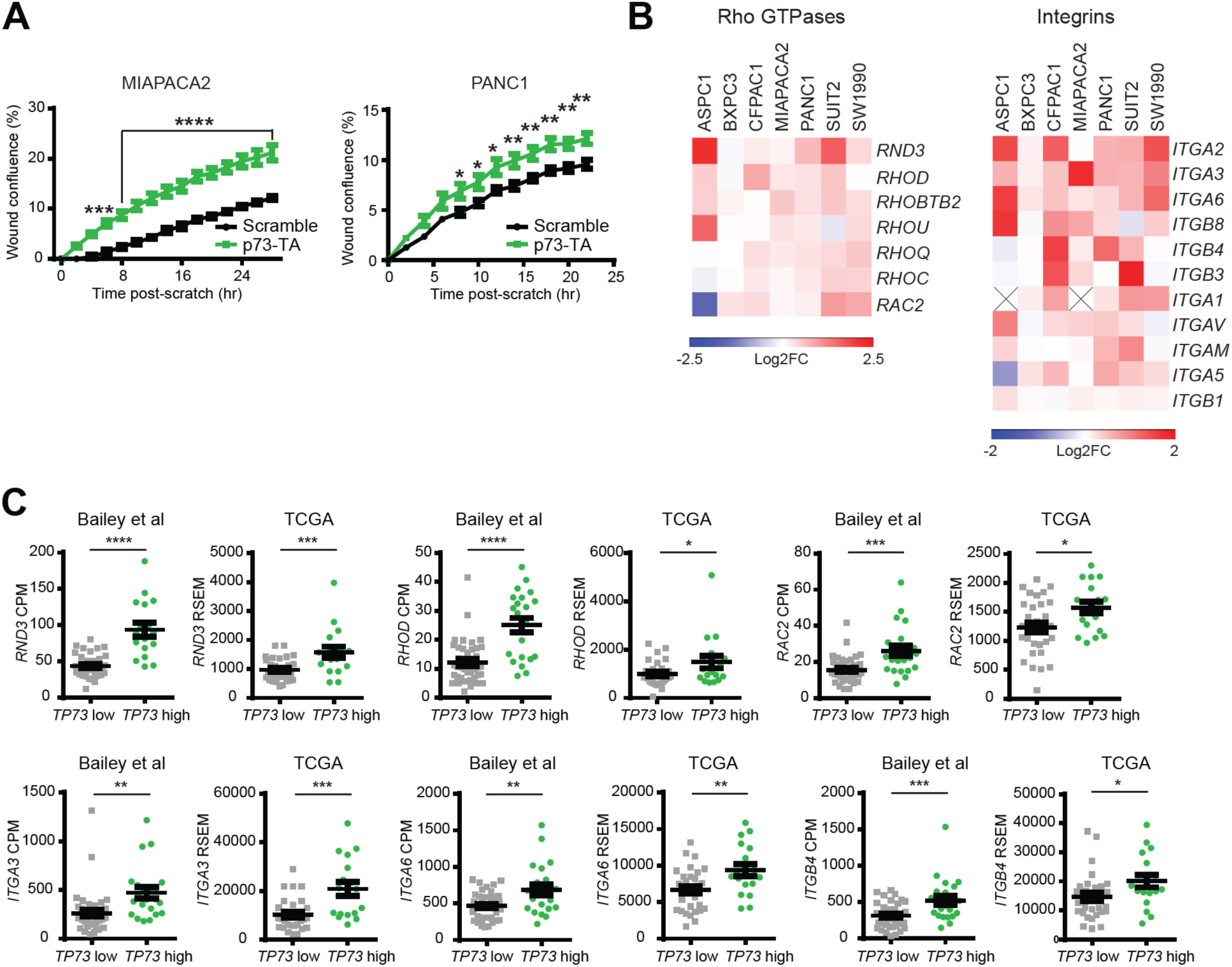
p73-TA promotes cell migration and invasion in PDAC cells and upregulates Rho GTPases and integrins. Related to Figure 4. (*A*) Scratch assays following activation of p73-TA compared to the control (Scramble). Quantification of % wound confluence plotted at the indicated time points post-scratch. P-values versus Scramble calculated using two-way ANOVA with Sidak’s multiple comparisons test. Mean ± SEM shown (n=3). *, P ≤ 0.05; **, P ≤ 0.01; ***, P ≤ 0.001; ****, P ≤ 0.0001. (*B*) RNA-seq analysis following p73-TA activation in a panel of PDAC cell lines. Heatmaps show log2 fold-change in the expression level of a subset of RhoGTPases and integrins that are significantly upregulated in two or more cell lines. *ITGA1* is expressed at low levels in ASPC1 and MIAPACA2 regardless of the p73-TA activation status, thus not depicted and marked as X. (*C*) A subset of RhoGTPases and integrins that are expressed at higher levels in *TP73*^high^ tumors compared to the *TP73*^low^ tumors in both of the indicated data sets. P-values calculated using unpaired t-test. Mean ± SEM shown. *, P ≤ 0.05; **, P ≤ 0.01; ***, P ≤ 0.001; ****, P ≤ 0.0001.

**Supplementary Figure 6.**
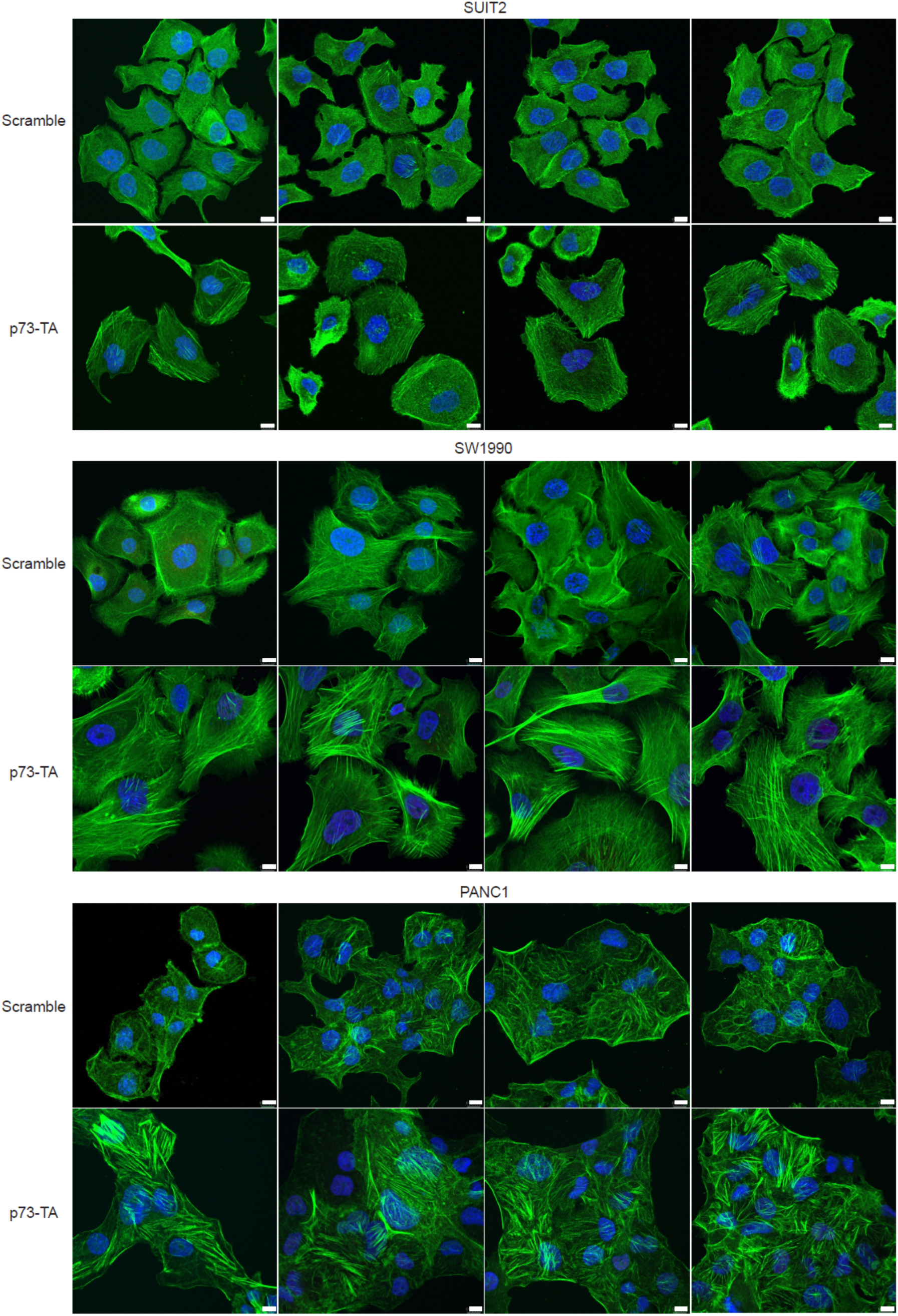
Phalloidin staining to visualize F-actin. F-actin (green) and nuclei (stained with DAPI, blue) shown in control (Scramble) and p73-TA activated cells (p73-TA). Scale bar indicates 10 µm. Related to Figure 4.

## Supplementary Information

Supplementary Table 1. Sequences of sgRNAs and RT-qPCR primers used in this study

Supplementary Table 2. List of genes depicted in the heatmap in Fig.2 C, in the same order as it appears on the heatmap.

## Materials and Methods

### Cell lines/organoids

Human PDAC cell lines ASPC1 (CRL-1682), BXPC3 (CRL-1687), MIAPACA2 (CRL-1420), SW1990 (CRL-2172), PANC1 (CRL-1469), CFPAC1 (CRL-1918), HS766T (HTB-134), and CAPAN2 (HTB-80) were purchased from ATCC. SUIT2 (JCRB1094) and KP2 (JCRB0181) were purchased from JCRB. PATU8988T (ACC-162), PATU8902 (ACC-179), and YAPC (ACC-382) were purchased from DSMZ. Human PDAC organoid hM1a (Tiriac et al. 2018) was gifted from the Tuveson Lab. ASPC1, BXPC2, MIAPACA2, SW1990, CFPAC1, HS766T, CAPAN2, SUIT2, KP2, and YAPC were cultured in RPMI 1640 (Corning, Cat# 10-040-CV) and PANC1, PATU8988T, PATU8902, and HEK293T were cultured in DMEM (Corning, Cat# 10-013-CV), all supplemented with 10% FBS and 1% Penicillin/Streptomycin (Thermo Fisher Scientific, Cat# 15140122). hM1a was cultured as described in (Boj et al 2015, Tiriac et al 2018). Briefly, organoids were grown in Matrigel (Corning, Cat# 354230) domes and supplemented with Human Complete Feeding Medium composed of: advanced DMEM/F12 (Life Technologies, Cat# 12634-028), HEPES 10mM (Life Technologies, Cat# 15630-130), Glutamax (Life Technologies, Cat# 35050-079), A83-01 500nM (Tocris Bioscience, Cat# 2939), hEGF 50ng/mL (PeproTech, Cat# AF-100-15), mNoggin 100ng/mL (PeproTech, Cat# 250-38), hFGF10 100ng/mL (PeproTech, Cat# 100-26), hGastrin I 0.01μM (Tocris Bioscience, Cat# 3006), N-acetylcysteine 1.25mM (Sigma-Aldrich, Cat# A9165-5G), Nicotinamide 10mM (Sigma-Aldrich, Cat# N0636-100G), PGE2 1μM (Tocris Bioscience, Cat# 2296), B27 supplement 1X final (Life Technologies, Cat# 17504044), R-spondin1 (Trevigen, Cat# 3710-001-K) conditioned media 10% final, Afamin/Wnt3A conditioned media (Osaka University) 50% final. Cell lines and organoids were grown at 37 °C, 5% CO_2_.

### Plasmid construction

To generate the LentiV-TP73-TA-neo vector, *TP73-TA* cDNA from HA-p73α-pcDNA2 (addgene, Cat# 22102) (Jost et al. 1997) was subcloned into the LentiV-neo empty vector (Somerville et al 2019). For CRISPR-based KO and competition-based proliferation assays, sgRNAs were subcloned into the LRNG vector (addgene, Cat# 125593). For CRISPR-based activation experiments, LentiV-dCas9-VPR-Blast vector was generated by replacing the Cas9 sequence from the lentiCas9-Blast (addgene, Cat# 52962) with the SP-dCas9-VP64-p65-Rta (VPR) sequence from the SP-dCas9-VPR vector (addgene, Cat# 63798) (Chavez et al. 2015); sgRNAs were subcloned into the pCRISPRia-v2 (addgene, Cat# 84832) (Horlbeck et al. 2016). sgRNA sequences are listed in Table S1.

### Lentivirus production and infection

Lentivirus was produced in HEK293T cells by transfecting plasmid DNA of interest with packaging plasmids (VSVG and psPAX2) using Polyethylenimine (PEI 25000; Polysciences, Cat# 23966-1). Media were replaced 6-8 hrs following transfection. Virus-containing supernatants were then collected every 24 hrs for 48 hrs and filtered through 0.45 μm filter prior to use.

To infect human PDAC cell lines, cell suspensions were mixed with virus-containing supernatants and centrifuged at 600 g for 30 minutes at room temperature. Following centrifugation, cells were plated in tissue culture plates, and virus-containing supernatants were replaced with fresh media after 24 hrs post-infection. To infect human PDAC organoids, virus-containing supernatants were first concentrated 10x using Lenti-X Concentrator (Takara, Cat# 631232). Single cell-suspended organoids were mixed with concentrated virus, supplemented with polybrene to a final concentration of 4 μg/ml, and centrifuged at 800 g for 2 hrs at room temperature. After centrifugation, virus-containing media were removed, and organoids were plated in Matrigel domes supplemented with fresh Human Complete Feeding Media. When selection was used to establish stable cell lines, corresponding antibiotics were added 48 hrs post-infection.

### CRISPR-based targeting

To generate cell lines in which p73 has been stably knocked out, cells were first infected with the LentiV-Cas9-puro vector (addgene, Cat# 108100) and selected with puromycin (3 μg/ml) to stably express Cas9. Subsequently, cells were infected with LRNG plasmids subcloned with p73 all isoform targeting (LRNG-TP73all), p73-TA-specific targeting (LRNG-TP73-TA), or control (LRNG-N) sgRNAs and selected with G418 (1 mg/ml). For competition-based proliferation assays, cells were infected as described above but without selection. % GFP was measured starting on day 3 post-sgRNA infection and then every two or three days until the end of experiment. GFP was measured using Guava easyCyte Flow Cytometers (Millipore).

To generate cell lines in which p73-TA has been stably activated, cells were first infected with the LentiV-dCas9-VPR-Blast and selected with blasticidin (10 μg/ml) to stably express catalytically-dead Cas9 (dCas9) fused with transcriptional activation domains VP64, p65, and Rta (together referred to as VPR) (Chavez et al. 2015). Subsequently, cells were infected with pCRISPRia-v2 plasmids subcloned with sgRNAs targeting upstream region of *TP73-TA* transcription start site (pCRISPRia-v2-TP73-TA) or scramble (pCRISPRia-v2-SC) sgRNAs and selected with puromycin (3 μg/ml).

### In vitro phenotypic assays

For cDNA overexpression experiments, cells were infected with LentiV-TP73-TA-neo or LentiV-neo empty vector as a control and selected with G418 (1 mg/ml). For CRISPR-based activation experiments, dCas9-VPR expressing cells were infected with pCRISPRia-v2-TP73-TA or pCRISPRia-v2-SC as a control and selected with puromycin (3 μg/ml). For CRISPR-based KO experiments, Cas9 expressing cells were infected with LRNG-TP73-TA, LRNG-TP73all, or LRNG-N as a control and selected with G418 (1 mg/ml). On day 7 post-infection, cells were counted by trypan blue exclusion and used for assays described below.

For cell proliferation assays, 2000 cells were plated in each well of 96-well plate (Corning, Cat# 3603) in triplicate. Number of viable cells on day 0 was measured at 6 hrs post-seeding. Subsequent measurements were taken every 24 hrs for 5 days using CellTiterGlo Luminescent Cell Viability Assay kit (Promega) and SpectraMax plate reader (Molecular Devices) following manufacturer’s instructions.

For scratch assays, cells were plated to confluency in wells of Incucyte Imagelock 96-well plate (Essen Bioscience, Cat# 4379) in triplicate. After cells have attached to the plate, media were replaced with serum-free media to start serum starvation. Following 18-24 hrs of serum starvation, scratch was made using Incucyte Woundmaker Tool (Essen Bioscience, Cat# 4563) according to manufacturer’s instructions. Wound closure was imaged every 2 hrs using Incucyte S3 Live-Cell Analysis System (Essen Bioscience). Relative wound density was quantified using Incucyte Scratch Wound Analysis Software Module (Essen Bioscience, Cat# 9600-0012).

3D Matrigel colony formation assays were performed as described previously (Somerville et al. 2018). Briefly, 5000 cells were resuspended in 1ml of RPMI supplemented with 5% Matrigel and 2% FBS and plated in each well of Costar 24-well Ultra-Low Attachment

Multiple Well Plates (Corning, Cat# 3473) in triplicate. Bright-field images were taken on day 7 post-seeding using 2x and 4x objectives. Colony number was quantified from two 2x images per well and colony size was quantified from four 4x images per well using ImageJ software.

Transwell assays were performed following Cell Invasion Assay protocol from Corning. Briefly, each insert of BioCoat Control Inserts with 8.0 μm PET Membrane (Corning, Cat# 354578) was coated with 100 μl of diluted Matrigel (final concentration of 300 μg/ml; diluted in buffer composed of 0.01M Tris pH 8.0 and 0.7% NaCl). Coated inserts were incubated at 37 °C, 5% CO_2_ for 2 hrs prior to plating the cells. Equal number of uncoated inserts were prepared. All experiments were set up in triplicate. 1×10^5^ cells were resuspended in 500 μl of serum-free media and added to each coated or uncoated insert, and 750 μl of 10% FBS media were added to the respective well of the plate as a chemoattractant. Cells were incubated for 20-24 hrs. After incubation, non-invading cells from the apical side of the inserts were removed using cotton swabs. Subsequently, inserts were fixed with 100% ice-cold methanol, stained with 0.1% Crystal violet solution, and air-dried for 24 hrs. Bright-field images were taken using 4x and 20x objectives: 4x images were used for figure representation; the number of invading or migrating cells were quantified from three 20x images per insert using ImageJ software. To control for differences in cell proliferation rate after migration/ invasion, data were represented as % Invasion = (the number of invading cells through Matrigel coated insert/ the number of migrating cells through uncoated insert) * 100.

### RNA extraction and RT-qPCR

Total RNA was extracted using TRIzol (Thermo Fisher Scientific, Cat# 15596018) and subjected to DNAase treatment (Thermo Fisher Scientific, Cat# EN0521) following manufacturer’s instructions. cDNA was synthesized using qScript cDNA SuperMix (Quanta bio, Cat# 95048-5000), followed by RT-qPCR using Power SYBR Green Master Mix (Thermo Fisher Scientific, Cat# 4368577) and primers in final concentration of 0.2μM on an ABI 7900HT fast real-time PCR system. Each sample was run in triplicate and 5ng of cDNA was used per reaction. Gene expression was normalized to *GAPDH*. Primers are listed in Table S1.

### Western blot analysis

To analyze stably knocked-out, activated, or overexpressed cells, cells were harvested on day 7 post-infection of sgRNAs or cDNA overexpression vectors. Viable cells were collected, washed with 1x PBS, and resuspended in 1x Laemmli Sample Buffer (2x Laemmli Sample Buffer diluted with 1x PBS, supplemented with 5% β-mercaptoethanol). Cells were then boiled for 30 min and centrifuged for 10 min at top speed. Supernatant was separated by the standard SDS-PAGE gel electrophoresis procedure, followed by transfer to nitrocellulose membrane and immunoblotting. For primary antibodies, p73 (Cell Signaling Technology, Cat# 14620) at 1:500, FOXJ1 (Atlas antibodies, Cat# HPA005714) at 1:700, and VINCULIN (Cell Signaling Technology, Cat# 4650) at 1:1000 were used. Proteins were detected using HRP-conjugated secondary antibodies and ECL substrate (Thermo scientific).

### Immunofluorescence staining

To stain cells in which p73-TA has been activated, cells on day 7 post-infection of sgRNAs were used. Cells were plated at 30-40% confluency on coverslips (Celltreat, Cat# 229172) placed in wells of 24-well plate. After cells have attached to the coverslips, media were replaced with serum-free media to start serum starvation. Following 18-24 hrs of serum starvation, cells were washed in 1x PBS, then fixed in 4% Paraformaldehyde (diluted in 1x PBS, Thermo Fisher Scientific, Cat# 50980494) for 15 min. Fixative was removed by washing the cells in 1x PBS three times. Cells were then permeabilized in 0.2% Triton X-100/1x PBS for 5 min followed by three washed in 1x PBS. Subsequently, cells were blocked in 3% -BSA/1x PBS for 30 min prior to incubation with fluorescence-conjugated Phalloidin (abcam, Cat# ab176753) at 1:000 for 1 hr at room temperature. Following three washes in 1x PBS, nuclei were counterstained with DAPI (Sigma Aldrich, Cat# D8417-10MG) for 10 min. Finally, cells were washed three times in dH2O and mounted with ProLong Diamond Antifade Mountant (Thermo Fisher Scientific, Cat# P36961). Phalloidin and DAPI staining were performed in dark. Images were acquired at 40x magnification using Leica SP8 confocal. To quantify % cells with actin polymerization, at least ten 40x images were analyzed per replicate in a total of three biological replicates.

### RNA-seq library construction

For RNA-seq experiments of cells in which p73 has been overexpressed, activated, or knocked-out, cells on day 7 post-infection of sgRNAs or overexpression vectors were used. RNA-seq library was constructed following the TruSeq RNA Sample Preparation v2 Guide (Illumina). Briefly, 1 μg of purified RNA was poly-A selected and fragmented using fragmentation enzyme. cDNA was synthesized using Super Script II (Thermo Fisher, Cat# 18064014), followed by end repair, 3’ adenylation, adapter ligation, and PCR enrichment of adapter-ligated DNA in the library. Library was quantified using Qubit Fluorometer (Thermo Fisher Scientific), and library size and purity was assessed using 2100 Bioanalyzer (Agilent). Equal molar quantities of libraries were multiplexed and sequenced on NextSeq platform with single-end reads of 50 bp. Approximately 30-50 million reads were obtained per sample.

### RNA-seq analysis

RNA-seq experiments of cDNA overexpression, CRISPR-based activation, and CRISPR-based KO were done in two biological replicates. Reads were mapped to hg38 genome using HISAT2 with default parameters (Kim et al 2015). Per gene mapped reads were then counted using HTSeq-count (Anders et al 2015) and custom gtf file containing only protein coding genes. Differentially expressed genes were analyzed using DESeq2 (Love et al. 2014) with default parameters. Log2 fold change values calculated by DEseq2 were used for all subsequent analysis below. To perform pre-ranked GSEA analysis (Subramanian et al 2005), genes were ranked by their log2 fold changes between control and experimental groups.

To compare gene expression changes between cDNA overexpression of p73-TAα and CRISPR-based activation of p73-TA, top 200 significantly upregulated genes in all three cell lines (ASPC1, CFPAC1, PANC1) in cDNA overexpressed samples were first identified. Genes with mean normalized counts of less than 1 in both control and experimental samples in at least one cell line were excluded from the analysis. Next, gene ontology analysis using Metascape with the Express analysis default parameters (Zhou et al., 2019) was performed, identifying *TP53*-pathway related terms such as PID P53 DOWNSTREAM PATHWAY (M145), GOBP APOPTOTIC SIGNALING PATHWAY (GO:0097190), GOBP EXTRINSIC APOPTOTIC SIGNALING PATHWAY (GO:0097191), and GOBP NEGATIVE REGULATION OF CELL POPULATION PROLIFERATION (GO:0008285) (MSigDB) (Liberzon et al 2011, Liberzon et al 2015). Finally, significantly upregulated genes that belong to the aforementioned terms were depicted in a heatmap as log2 fold changes in expression levels following overexpression of p73-TA.

To interrogate pathways enriched in CRISPR-based activation of p73-TA, top 500 significantly upregulated genes in two or more cell lines (out of seven cell lines) were first identified. Then, Metascape with the Express analysis default parameters was used to perform gene ontology analysis (Zhou et al., 2019).

To examine specific Rho GTPase and integrin family genes upon CRISPR-based activation of p73-TA, only genes that were significantly upregulated in two or more cell lines (out of seven cell lines) were included in the analysis. Log2 fold changes in expression levels following overexpression of p73-TA were depicted in heatmap.

Heatmaps were generated using Excel or Morpheus (Broad Institute, https://software.broadinstitute.org/morpheus).

### ChIP

For ChIP-seq experiments of cells in which p73-TA has been activated, cells on day 7 post-infection of sgRNAs were used. 5-10 million cells were used for H3K27ac ChIP-seq and 60-100 million cells were used for p73 ChIP-seq. Cells were harvested as single cell suspensions and crosslinked in 1% formaldehyde for 10 min at room temperature and then quenched with 0.125M glycine for 10 min at room temperature. Followed by a wash with ice-cold 1x PBS, cells were lysed in cell lysis buffer (10mM Tris-HCl pH 8.0, 10mM NaCl, 0.2% NP-40) supplemented with protease inhibitor for 15 min on ice. Nuclei were isolated by centrifugation at 3100 g for 5 min at 4°C. Next, for every 10 million cells, nuclei were lysed in 1 ml of nuclei lysis buffer (50mM Tris-HCl pH 8.0, 10mM EDTA, 1% SDS) supplemented with protease inhibitor for 10 min on ice. Using the 15 ml Bioruptor Pico tubes and sonication beads (diagenode) and Bioruptor Pico sonication device (diagenode), 1 ml of lysed nuclei was then sonicated in one 15 ml tube for 10 cycles at 30 sec on/off setting. Cell debris were pelleted by centrifugation at top speed for 10 min at 4°C and supernatant was kept to proceed with immunoprecipitation (IP). Prior to IP, supernatant was diluted in IP dilution buffer (20 mM Tris-HCl pH 8.0, 2 mM EDTA, 150 mM NaCl, 0.01% SDS, 1% Triton X-100) by 8 fold. For H3K27ac ChIP, sonicated chromatin was incubated with 2 μg of H3K27ac antibody (abcam, Cat# ab4729) and 25 μl of Dynabeads Protein A (Thermo Fisher Scientific, Cat# 10002D). For p73 ChIP, sonicated chromatin was incubated with 14 μg of p73-TA antibody (abcam, Cat# ab14430) and 100 μl of Dynabeads Protein A. IP was performed at 4°C overnight. Following IP, beads bound to chromatin were washed once with IP-wash 1 buffer (20 mM Tris-HCl pH 8.0, 2 mM EDTA, 50 mM NaCl, 0.1% SDS, 1% Triton X-100), twice with High-salt buffer (20 mM Tris-HCl pH 8.0, 2 mM EDTA, 500 mM NaCl, 0.01% SDS, 1% Triton X-100), once with IP-wash 2 buffer (10 mM Tris-HCl pH 8.0, 1 mM EDTA, 250 mM LiCl, 1% NP-40, 1% sodium deoxycholate), and twice with TE buffer (10 mM Tris-HCl, 1 mM EDTA, pH 8.0). Next, DNA was eluted from beads in 200 μl of nuclei lysis buffer at 65°C for 15 min with shaking. 12 μl of 5M NaCl and 2 μl of RNaseA (1mg/ml) were then added to the eluted DNA and incubated overnight at 65°C to reverse cross-link. DNA was treated with proteinase K at 42°C for 2 hrs and purified using MinElute PCR Purification Kit (Qiagen, Cat# 28004).

### ChIP-seq library construction

ChIP-seq library was constructed following TruSeq ChIP Sample Preparation Guide (Illumina). Briefly, ChIP DNA was end repaired, followed by 3’ adenylation, adapter ligation, and size selection by gel electrophoresis on 2% agarose gel. Then, 15 PCR cycles were used for final enrichment of adapter-ligated DNA in the library. Library was quantified using Qubit Fluorometer, and library size and purity was assessed using High Sensitivity chip (Agilent) on 2100 Bioanalyzer. Equal molar quantities of libraries were multiplexed and sequenced on NextSeq platform with single-end reads of 50 bp. Approximately 50-70 million reads were obtained per sample.

### ChIP-seq analysis

Reads were mapped to hg38 using Bowtie2 with default parameters (Langmead and Salzberg, 2013). MACS2 (Zhang et al 2008, Feng et al 2012) was then used to call peaks using input genomic DNA as a control, with the narrow peak and broad peak options for analyzing the H3K27ac and p73 ChIP-seq data sets, respectively. To compare H3K27ac peak profiles between control and experimental groups, all called peaks across the control and experimental groups were merged using BEDTools (Quinlan and Hall, 2010). This led to a union of all 161734 H3K27ac peaks. The squamous elements were lifted to hg38 from hg19 for the analysis (Somervaille et al. 2018). 1333 random elements from all H3K27ac peaks were chosen as controls. ChIP-seq peak intensities of the control or experimental data sets were then recounted and normalized at the squamous elements and the random elements using bamToGFF script (https://github.com/BradnerLab/pipeline). The average of H3K27ac peak intensities at regions of interest was then used to generate metagene plots with +/- 10kb extension around peak center. To visualize individual ChIP-seq tracks, BigWig files were uploaded to the UCSC genome browser.

### Publicly available datasets analysis

Publicly available patient datasets TCGA-PAAD and QCMG-PAAD (Bailey et al. 2016) were obtained from CBioPortal (Cerami et al 2012) and GSE71729 from Moffitt et al. (Moffitt et al 2015). All patient data sets were downloaded in January 2018. CCLE expression data set CCLE_RNAseq_rsem_transcripts_tpm_20180929.txt.gz (Ghandi et al 2019) was downloaded from www.broadinstitute.org/ccle. Normal tissue expression of *TP73* was downloaded from GTEx portal on 12/08/19 (The GTEx Consortium, 2015). *TP73* log2 (TPM+1) Expression Public 21Q1 and *TP73* Gene Effect (CERES) CRISPR (Avana) Public 21Q1 were obtained from depmap portal (DepMap, 2020) (Meyers et al 2017 and Dempster et al 2019, Ghandi et al 2019). Human PDAC organoid expression data set (Tiriac et al 2018) was provided by Tuveson lab. For survival analysis, all samples from Moffitt et al. and Bailey et al. were used, and only high purity samples from TCGA-PAAD were used. *TP73* mutational analysis was done in cBioPortal (Cerami et al 2012). To analyze % isoform expression of *TP73* in CCLE and human organoid data sets, samples with *TP73* tpm values less than 1 were excluded from the analysis. To analyze human TF expression in squamous and progenitor classes of tumors, squamous or progenitor samples from Bailey (n=55 samples) and high purity, Bailey cluster squamous or progenitor samples from TCGA-PAAD (n=65 samples) were included in the analysis. Human TFs described in Lambert et al. (Lambert et al 2018) with average expression of all samples greater than RSEM 20 (TCGA-PAAD; n=1236 TFs) and CPM 1 (Bailey et al; n=1229 TFs) were included in the analysis. To analyze human TF expression in *TP73* high and *TP73* low classes of tumors, all samples from Bailey et al. were used, and only high purity samples from TCGA-PAAD were used. Samples were designated as *TP73* high or low based on z-score expression values of > 0.2 or < -0.4, respectively. Here, human TFs described in Lambert et al. (Lambert et al 2018) with average expression of all samples greater than RSEM 20 (TCGA-PAAD; n=1301 TFs) and CPM 1 (Bailey et al; n=1249 TFs) were included in the analysis. Venn diagrams were created using BioVenn (Hulsen et al., 2008).

### Statistical analysis

GraphPad Prism was used for statistical analysis. P values from unpaired two-tailed Student’s t test or ANOVA were used to evaluate statistical significance, as indicated in the figure legends. Log-rank (Mantel-Cox) test was used to assess the median survival values and p-values of the Kaplan-Meier survival curves. ns, P > 0.05; *, P ≤ 0.05; **, P ≤ 0.01; ***, P ≤ 0.001; ****, P ≤ 0.0001.

